# Impact of Reduced Chlorophyll Levels in Leaves on Soybean Yield, Seed Composition, Pod/Seed Photosynthesis, and Chlorophyll Levels in Pod and Seed Tissues

**DOI:** 10.64898/2026.08.14.744892

**Authors:** Sarah I. Jones, Samantha S. Stutz, Eray Atalay, Yu Wang, Donald R. Ort, Young B. Cho

## Abstract

Soybean, a widely cultivated leguminous crop valued for its protein, amino acids, and oil, faces the challenge of maintaining protein levels, which have an inverse correlation with yield. Reducing leaf chlorophyll levels could increase seed protein levels without compromising yield; however, this is yet to be tested. Therefore, to understand the impacts of low chlorophyll mutations on soybean yield and seed composition, we screened and compared 25 low chlorophyll soybean mutants to their 11 dark green parents. PI548210 (Lincoln mutant) demonstrates a higher concentration of protein without affecting yield compared to its dark green parent PI548362 (Lincoln), suggesting it as a good candidate for further large-scale field trials. PI547555 (*Y11/y11,* Clark mutant) demonstrates a lower concentration of oil without impacting yield, alongside lower gross photosynthesis, but with chlorophyll levels in the pod and seed tissues that are comparable to its dark green parent PI548533 (Clark). These findings are consistent with the oil concentration of the soybean being influenced by pod and seed photosynthesis, which is correlated with pod height and row spacing. Chlorophyll levels in the leaf do not necessarily correlate with those in the pod and seed of low chlorophyll mutants, possibly due to substantially lower expression of chlorophyll synthesis genes in the pod and seed.

**SIGNIFICANCE:**

- PI548210 (Lincoln mutant), one of twenty-five low chlorophyll soybean mutants, demonstrates a higher concentration of soybean protein without affecting yield compared to its dark green parent (Figure 1 and Table 1).
- PI547555 (*Y11/y11*, Clark mutant), a low chlorophyll soybean mutant, demonstrates a reduced concentration of soybean oil without impacting yield, alongside lower gross photosynthesis in pod and seed tissues compared to its dark green parent (Figures 3 and Table 2). These findings suggest that the oil concentration of the soybean is influenced by pod and seed photosynthesis, which is in turn influenced by pod height and row spacing (Figure 2).
- Chlorophyll levels in the leaf do not necessarily correlate with those in the pod and seed of low chlorophyll mutants, possibly due to substantially lower expression of chlorophyll synthesis genes in the pod and seed (Figure 5-6).

## INTRODUCTION

Legumes contribute a significant share to the diets of humans, poultry, and livestock (Mandal and Mandal, 2000). Soybean is the most widely cultivated leguminous crop due to its high yield of protein, essential amino acids, and oil that is used especially for animal feed (Warrington et al., 2015). Soybean meal, which is composed mainly of crude protein, constitutes about 60% of soybean value while the production of oil makes up the remaining 40% (Pettersson and Pontoppidan, 2013). Because of its high protein content, soybean meal has been the preferred source of protein for the livestock and poultry industry for several decades. Soybean meal generally provides more protein, crude fiber, and amino acids tryptophan, threonine, isoleucine, and valine than meal from sorghum (*Sorghum bicolor*), corn (*Zea mays*), and other cereals. The improvement of levels of protein and amino acids in soybean would raise the crop’s economic value and contribute to the entire supply chain from growers to processors to consumers. However, improving levels of protein and amino acids in soybean is challenging due to protein’s inverse correlation with oil content and seed yield (Chung et al., 2003; Chaudhary et al., 2015; Bandillo et al., 2015; Kim et al., 2016; Patil et al., 2017).

**Figure 1.**
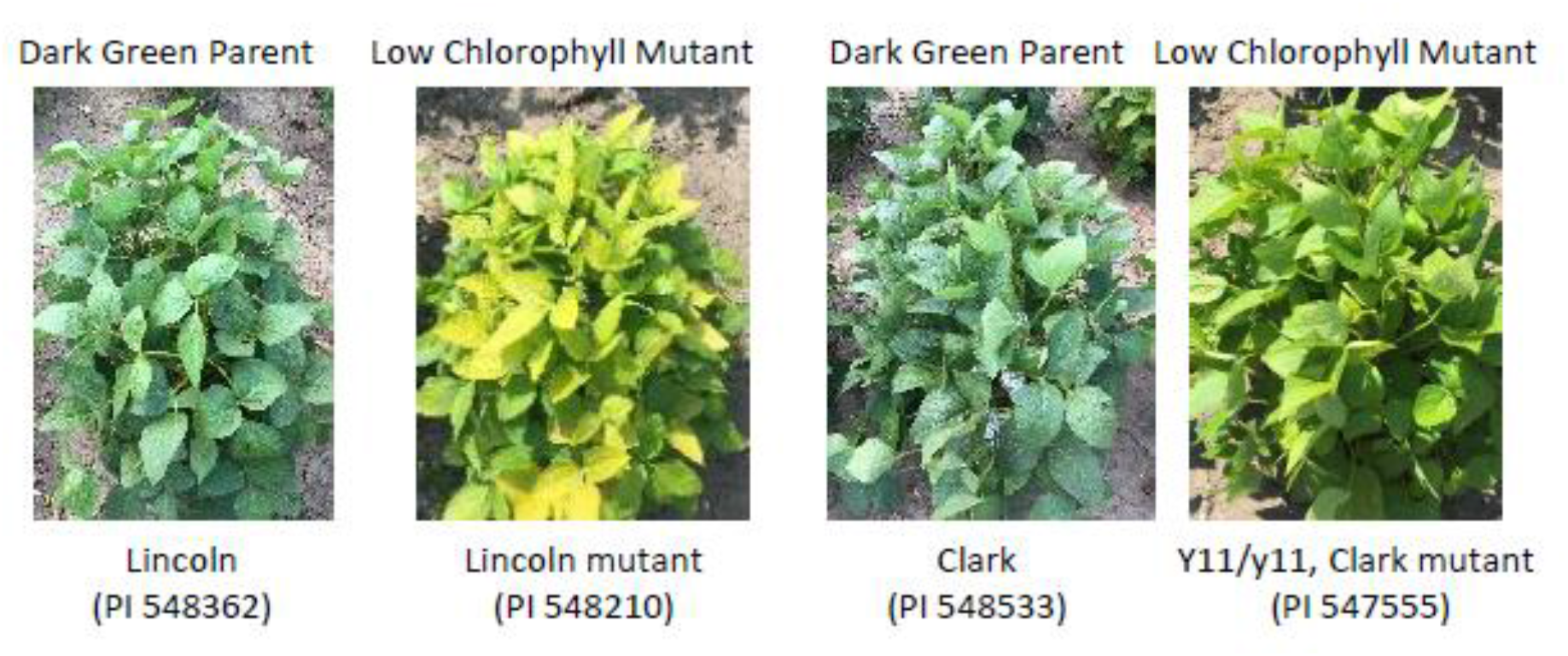
Two low chlorophyll mutants are as healthy as their dark green parents. Lincoln and its low chlorophyll mutant, left; Clark and its low chlorophyll mutant, known as Y11/y11, right. It can be seen by eye that the plants have low chlorophyll (light green/yellow leaves) but a similar growth habit to their dark green parents. See Supplemental Figures 1-4 for contrast, where low chlorophyll mutants are stunted in growth compared to their dark green parents.

**Figure 2.**
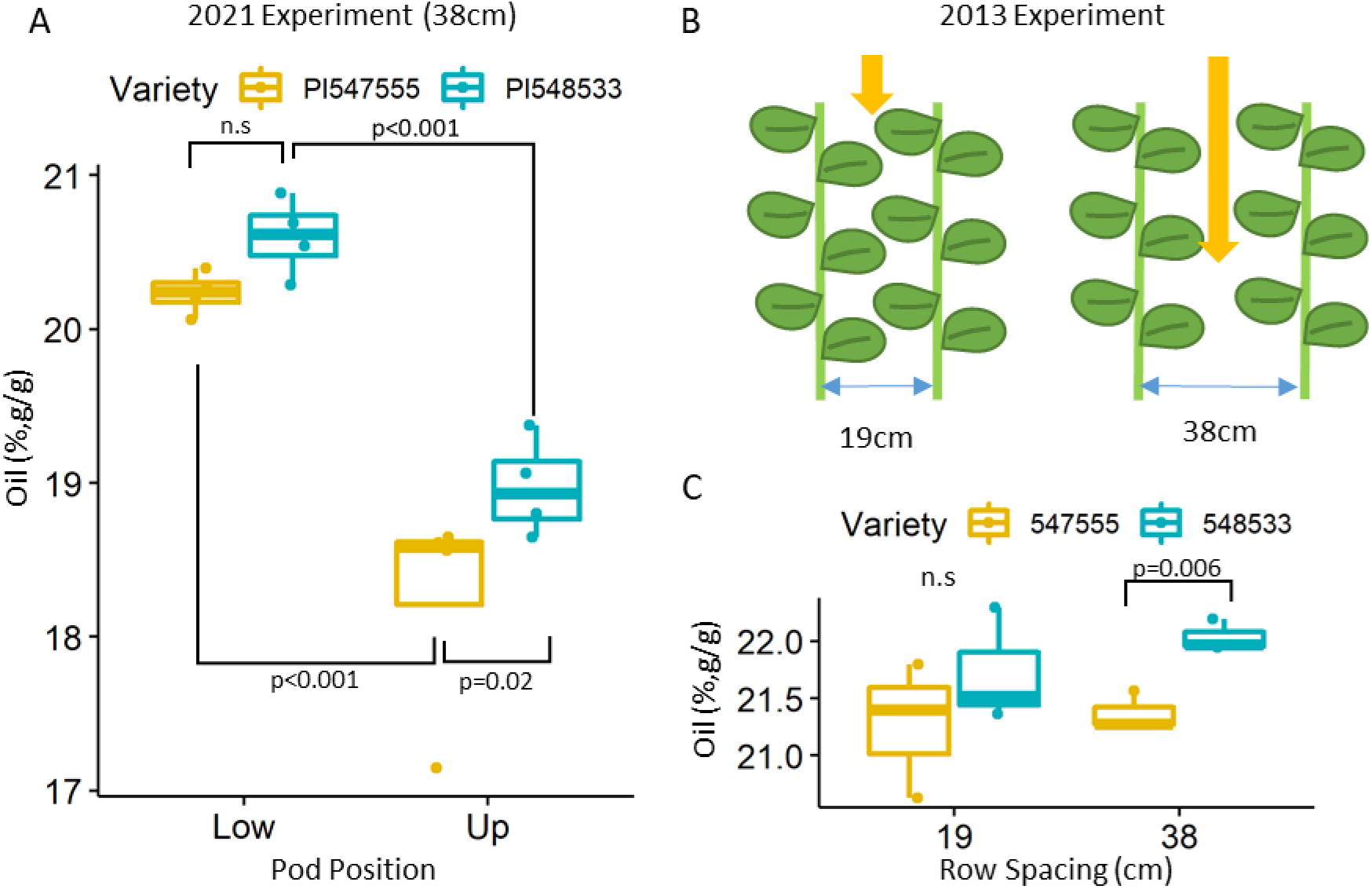
Low chlorophyll mutant (Y11/y11, PI547555) and its parent (Clark, PI548533) differ in concentration of seed oil, which interacts with height of pod and row spacing. The box plots show the median (central line), the lower and upper quartiles (box) and the minimum and maximum values (whiskers). The statistical analysis was done using ANOVA with linear mixed model (n=3 blocks, alpha=0.05). Least squares mean is used to compare. N.s., non- significant in the analysis. A. Concentration of oil in low chlorophyll mutant seeds from the upper canopy decreased by 4% compared to the dark green parent (18.2% vs 19%) while there was no difference between them in the seeds from the lower canopy (20.2% vs 20.6%). B. Schematic layout of 2013 field setting showing two different row spacings. C. Concentration of oil in low chlorophyll mutant decreased by 2% in 38cm spacing (21.4% vs 22%) while there was no difference in 19cm spacing (21.3% vs 21.7%) in 2013 field.

**Figure 3.**
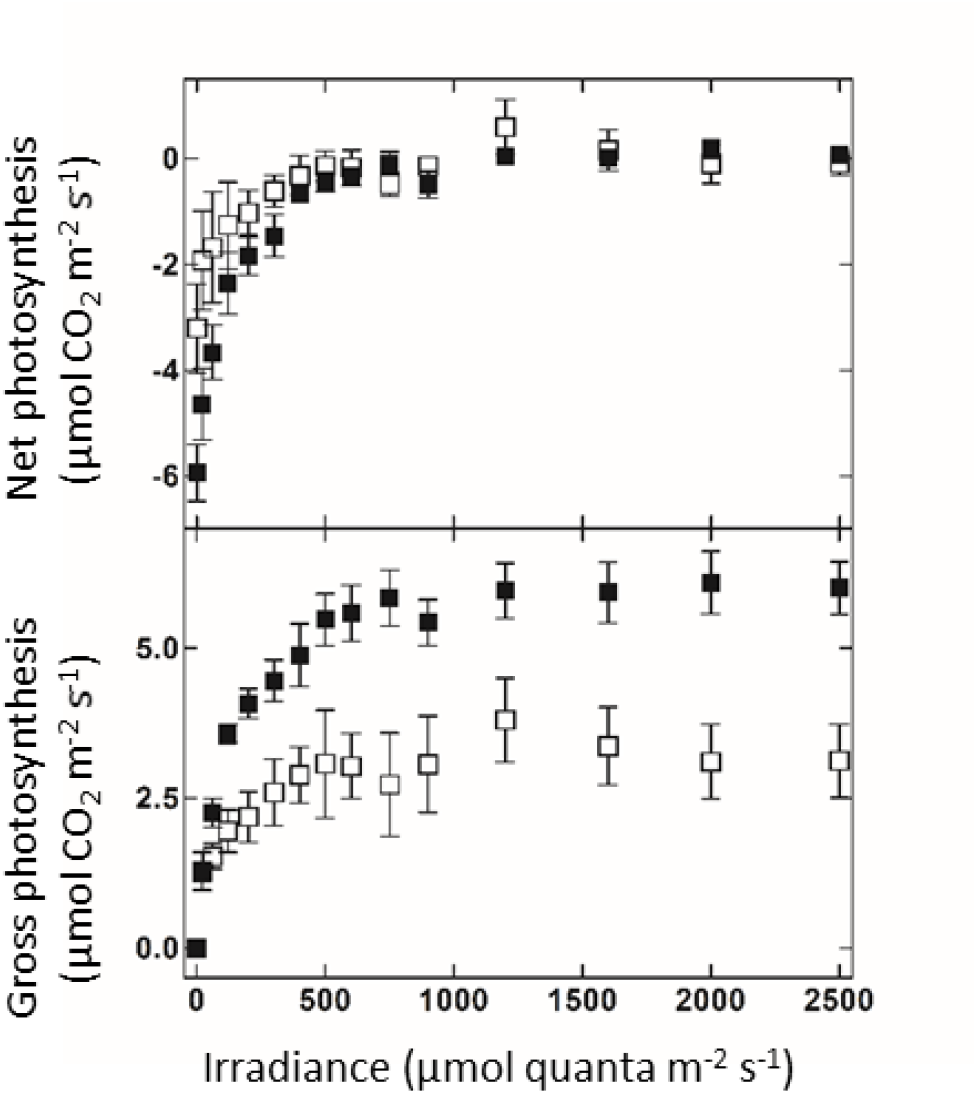
Light response curve of low chlorophyll mutant (*Y11/y11*, PI547555) and its parent (Clark, PI548533). Rates of net and gross photosynthesis of low chlorophyll (white) and dark green parents (black) pods under field conditions. Each dot represents a value (n=4) ±SE. We assumed that the seeds greatly inhibited the transmittance of light through the pod and used photosynthetic photon flux density for a single-side.

**Figure 4.**
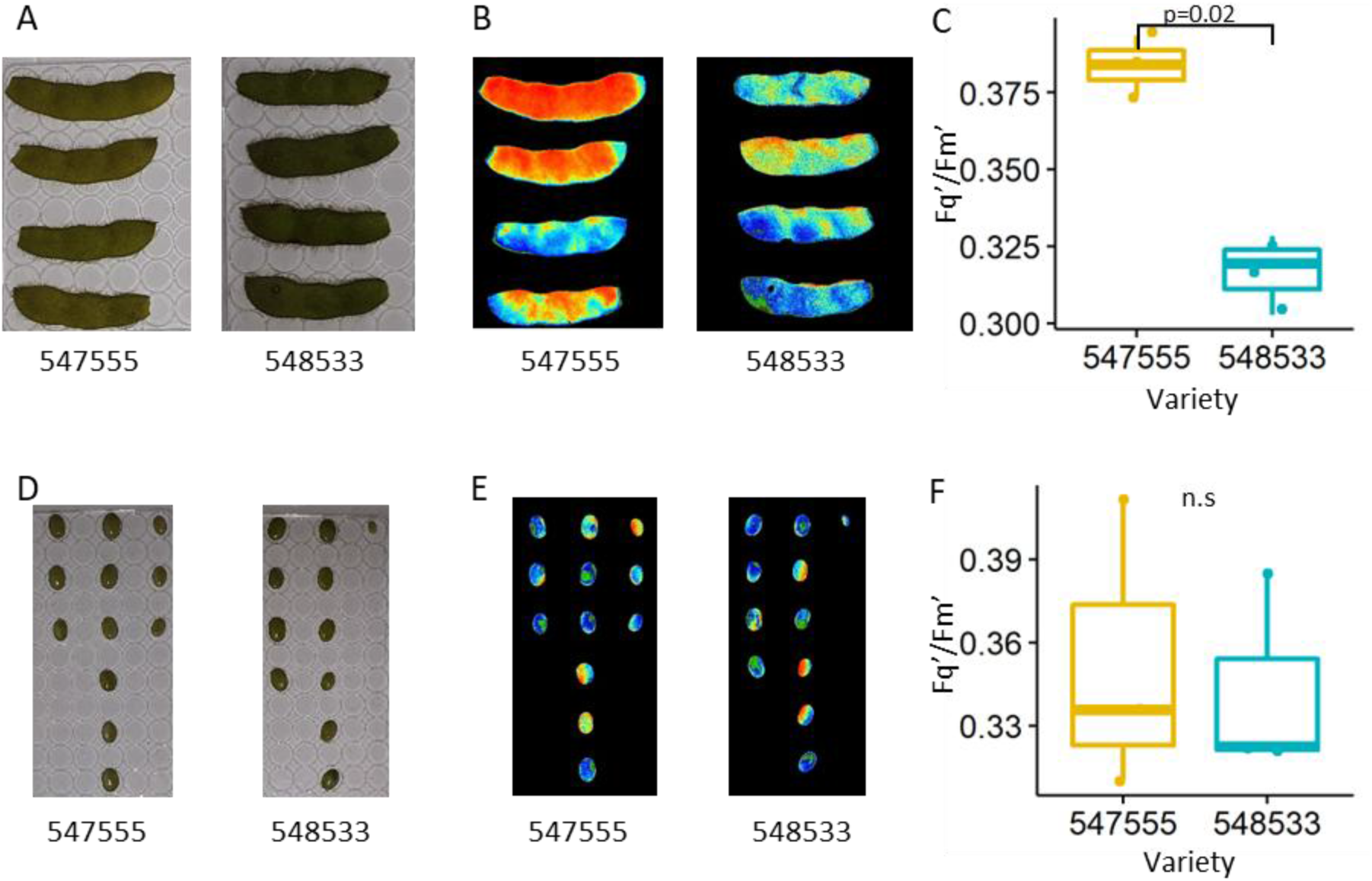
Low chlorophyll mutant (Y11/y11, PI547555) and its parent (Clark, PI548533) differ in chlorophyll fluorescence in immature pods, but not seeds. Operating efficiency of Photosystem II (Fq’/Fm’) of low chlorophyll mutant (*Y11/y11*, PI547555) and dark green parent (Clark, PI548533) from 2021 field. The statistical analysis was done using ANOVA with linear mixed model (n=3 blocks, alpha=0.05). N.s., non-significant in the analysis. A and D. Pictures of immature pods and seeds. Seeds are 100-200mg fresh weight. B and E. Chlorophyll fluorescence images of the same samples. C and F. Fq’/Fm’ in pods of low chlorophyll mutant is higher (0.384) than in dark green parent (0.317) (C), while Fq’/Fm’ in seeds shows no difference (F).

**Figure 5.**
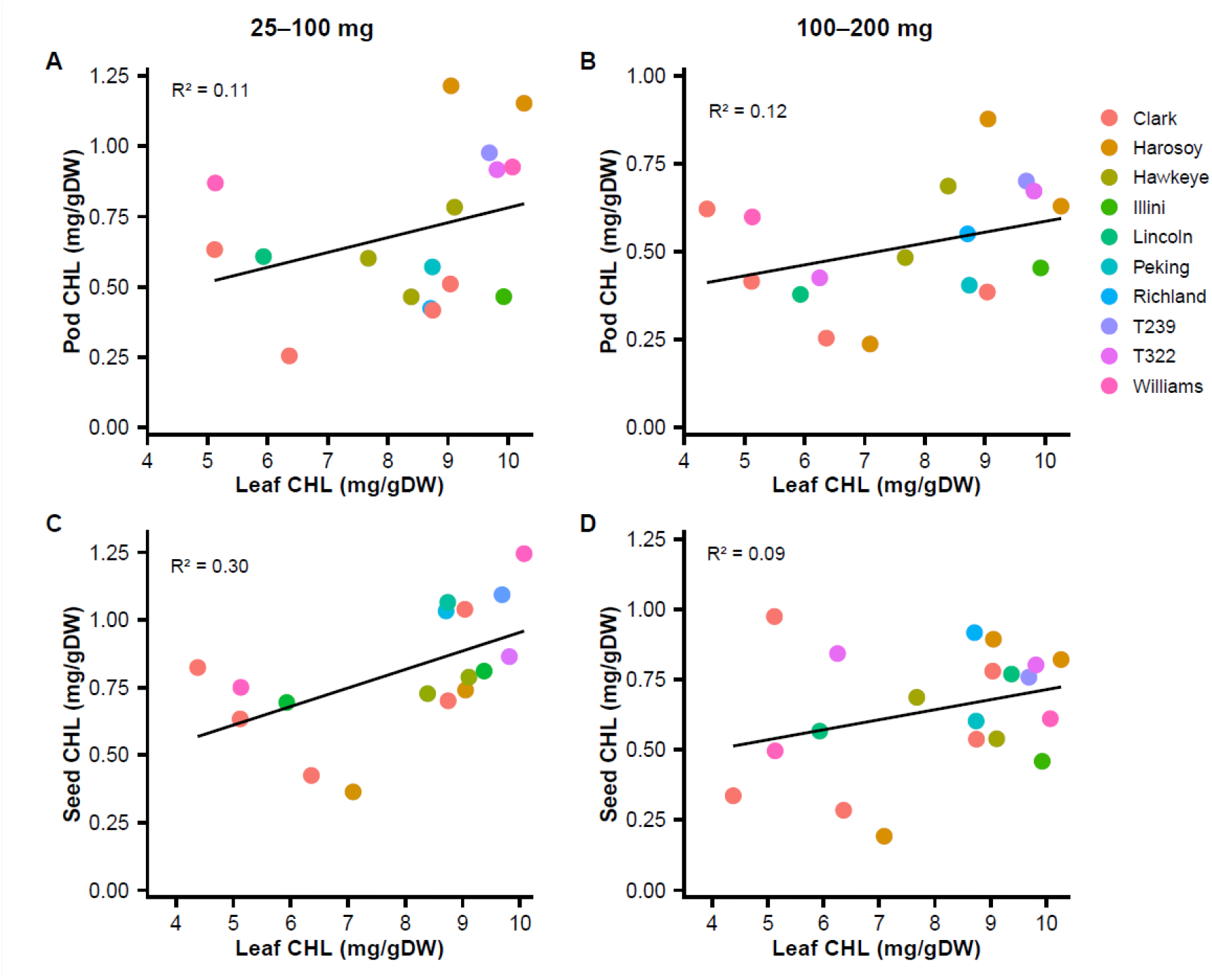
(greenhouse). Correlation between the level of leaf chlorophyll (x-axis: SPAD reading) and the level of immature pod or seed chlorophyll (y-axis, mg/g DW). Line represents the linear regression model. R-squared is a coefficient of determination, the percentage of the response variable variation that is explained by the linear model. Pod is labeled by the fresh weight of seeds it contained. A. Level of chlorophyll of 25-100mg pod (n=18). B. Level of chlorophyll of 100-200mg pod (n=17) . C. Level of chlorophyll of 25-100mg seed (n=17). D. Level of chlorophyll of 100-200mg seed (n=20).

**Figure 6.**
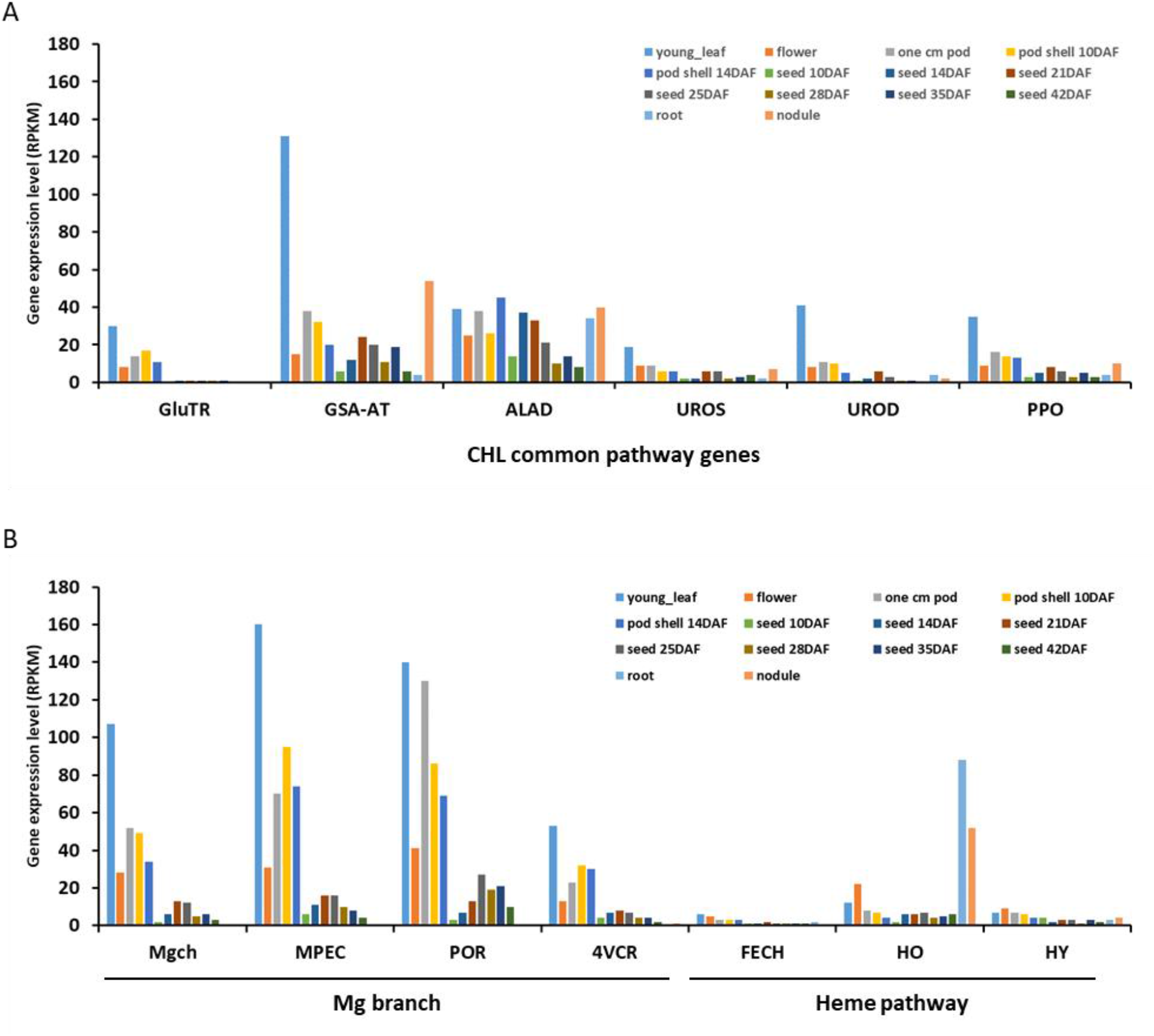
Levels of gene expression in chlorophyll synthesis pathway. A. CHL common pathway genes; Glutamyl-tRNA reductase (GluTR). Glutamate 1- semialdehyde aminotransferase (GSA-AT). ALA dehydratase (ALAD). Uroporphyrinogen III synthase (UROS). Uroporphyrinogen III decarboxylase (UROD). Protoporphyrinogen IX oxidase (PPO). B. Mg branch; Mg-chelatase (Mgch). Magnesium-protoporphyrin IX monomethyl ester cyclase (MPEC). Protochlorophyllide reductase (POR). 3,8-divinyl protochlorophyllide a 8-vinyl-reductase (4VCR). Heme pathway; Ferrochelatase (FECH). Heme oxygenase (HO). Phytochromobilin synthase (HY). Data come from Severin et al (2010). RPKM, reads per kilobase per million mapped reads. DAF, days after flowering. The source seed is experimental line A81-356022 which was generated by introgressing G. soja (PI468916) into G. max (A81-356022).

**Table 1.** Comparison of seed yield, weight, seed composition between low chlorophyll mutants and their dark green parents. ANOVA is used with linear mixed model (random effect = block, fixed effect = variety). Least squares mean is used to compare. For yield and seed composition, N=4 blocks. For leaf chlorophyll (SPAD), N=40. Yield is average yield per plant (g). n.s. = not significant.

| Variety | Yield<br>(g) | Seed<br>weight<br>(g/100<br>seeds) | Leaf CHL<br>(mg/gDW) | Concentration (g/g, %) |  |  |  |  |  |
| --- | --- | --- | --- | --- | --- | --- | --- | --- | --- |
|  |  |  |  | Oil | Protein | Lys | Cys | Thr | Met |
| Lincoln (PI 548362) | 32.3 | 15.0 | 9.4 | 20.9 | 42.6 | 2.5 | 0.5 | 1.5 | 0.5 |
| Lincoln mutant (PI 548210) | 30.9 | 11.8 | 3.6 | 17.4 | 44.9 | 2.6 | 0.5 | 1.5 | 0.5 |
| p-value | n.s. | 0.01 |  | <0.001 | 0.009 | n.s. | 0.03 | n.s. | 0.05 |
| Difference (%) |  | -21.3 |  | -16.7 | 5.4 |  | 5.2 |  | 3.6 |
| Clark (PI 548533) | 41.6 | 16.5 | 9.1 | 19.8 | 43.5 | 2.6 | 0.5 | 1.5 | 0.5 |
| Y11/y11 (PI 547555) | 39.3 | 15.1 | 4.4 | 19.2 | 43.7 | 2.5 | 0.5 | 1.5 | 0.5 |
| p-value | n.s. | 0.02 |  | 0.03 | n.s. | n.s. | n.s. | n.s. | n.s. |
| Difference (%) |  | -8.5 |  | -3.0 |  |  |  |  |  |

**Table 2.** Pod photosynthetic parameters for low chlorophyll mutant (Y11/y11, PI547555) and its parent (Clark, PI548533). Photosynthesis was measured 1 September through 15 September 2021 at the University of Illinois Energy Farm in Urbana, IL, USA. The statistical analysis was done using ANOVA with linear mixed model (alpha=0.05). N=4 ± SEM for Clark and N=3 ± SEM for Y11.

| ($\mu\text{mol CO}_2 \text{ m}^{-2} \text{ s}^{-1}$) | Clark (PI 548533) | Y11/y11 (PI 547555) | p-value |
| --- | --- | --- | --- |
| Respiration at 0 $\mu\text{mol m}^{-2} \text{ s}^{-1}$ | $9.0 \pm 0.8$ | $6.7 \pm 1.7$ | |
| Gross photosynthesis at 2000 $\mu\text{mol m}^{-2} \text{ s}^{-1}$ | $9.2 \pm 0.8$ | $6.5 \pm 1.0$ | |
| Gross photosynthesis at 300 $\mu\text{mol m}^{-2} \text{ s}^{-1}$ | $6.7 \pm 0.3$ | $5.3 \pm 1.0$ | |
| Net photosynthesis at 2000 $\mu\text{mol m}^{-2} \text{ s}^{-1}$ | $0.3 \pm 0.3$ | $-0.2 \pm 0.9$ | |
| Net photosynthesis at 300 $\mu\text{mol m}^{-2} \text{ s}^{-1}$ | $-2.3 \pm 0.7$ | $-1.4 \pm 0.7$ | |

The balance between carbon (C, ultimately leads to oil) and nitrogen (N, ultimately leads to protein) has a strong influence on seed composition. The N uptake during early seed development has a substantial impact on the protein composition of soybean seeds (Krishnan, 2001; Paek et al., 1997). The well-known inverse correlation between soybean seed protein concentration and both yield and oil content is thought to result from tightly linked QTLs or pleiotropic genetic effects that compete for carbon and nitrogen allocation (Diers et al., 1992; Nichols et al., 2006; Pathan et al., 2013; Sebolt et al., 2000; Munier-Jolain and Salon, 2005). One example of this competition is that leaves can perform either C assimilation or N remobilization, which are mutually exclusive processes (Havé et al., 2017). Leaves assimilate C and export it in the form of sugars to the seeds, but during senescence, lower photosynthetic activity reduces C assimilation and sugar export, ultimately reducing oil in seeds. In contrast, during senescence, N is transferred from leaves to seeds, following increased protein breakdown in leaves (Masclaux-Daubresse et al., 2008). Because of these differences, delaying senescence has opposing effects. Maintaining green leaves continues C assimilation, leading to increased seed yield and oil (Havé et al., 2017); however, it also delays N remobilization, often resulting in decreased seed N (and thus protein) content (Havé et al., 2017).

Plants significantly overinvest N in chlorophyll biosynthesis (Li et al., 2013; Slattery et al., 2017; Kirst et al., 2017; Gu et al., 2017; Sakowska et al., 2018). Chlorophyll is the most dominant tetrapyrrole in higher plants, making up ∼14% of the leaf’s N budget (Evans and Clarke, 2019). Modeling studies predicted about 9% of N can be saved by reducing chlorophyll (Song et al., 2017; Walker et al., 2018). It was proposed that saved N from lowering chlorophyll can increase seed N (and thus protein) without penalty on yield because more free N will be available in the seed filling stages without compromising overall canopy photosynthesis, thus increasing N content in seeds without decreasing the C content (and thus oil), as demonstrated by a recent study (Cho et al., 2024).

Organisms that have naturally invested less in chlorophyll biosynthesis (low chlorophyll mutants) are abundant in germplasm collections of algae and plants, due in part to easy phenotyping. Low chlorophyll mutants sometimes perform better than dark green relatives (reviewed in Slattery and Ort, 2021). Low chlorophyll mutants in both *Chlamydomonas* (truncated light antennae 1*, tla1*) and cyanobacteria (phycobilisome light-harvesting antenna, *pla*) increase light penetration and photosynthetic efficiency in cultures (Melis, 1999; Polle et al., 2002; Mitra and Melis, 2008; Kirst et al., 2014). Low-chlorophyll rice plants show noticeably higher yields than the dark green parent at high planting densities because their canopy allows for more light penetration, offsetting some of the over-saturation at the top of the canopy (Li et al., 2013; Gu et al., 2017). In soy, a heterozygous low chlorophyll *Y11/y11* mutant (PI547555) shows similar or higher rates of canopy photosynthesis (per unit leaf area) than its dark green Clark parent (PI548533), despite more than 50% reduction of the leaf chlorophyll concentration (Pettigrew et al., 1989; Slattery et al., 2017). However, the effect of low leaf chlorophyll on seed composition has not been explored yet.

A second factor that can influence seed composition is pod position under the canopy and other canopy structure changes, which can create differing assimilate availability or micro- environments. Poeta et al. (2016) reported that high-protein genotypes often have reduced leaf area and harvest index (percentage of biomass devoted to seeds as opposed to other tissues) compared to high-yielding genotypes, reinforcing the trade-off between protein and yield; the smaller of these high-protein seeds were associated with less C per seed but more canopy biomass production. Canopy position can also have an effect: seeds at the top of the canopy contain more protein and less oil than seeds at the bottom of the canopy (Collins & Cartter, 1956), a positional effect repeatedly observed in both low- and high-protein breeding lines (Escalante & Wilcox, 1993a; Escalante & Wilcox, 1993b). More recently, Huber et al. (2016) also reported that seeds produced at the top of the canopy have higher concentrations of protein and less oil, and additionally, lower concentrations of Mg, Fe, and Cu compared to seeds produced at the bottom of the canopy. Environmental factors such as temperature, irradiance, light quality, and humidity affect soybean seed composition, all of which can differ in micro- environments from top to bottom under the canopy (Wolf et al., 1982; Carrera et al., 2009; Rotundo and Westgate, 2009; Carrera et al., 2011; Baldocchi et al., 1983). The micro- environment under the canopy is changed when neighboring plants are removed, leading to higher protein and less oil (Huber et al., 2016).

One potential mechanism by which canopy structure influences seed composition is through pod and seed photosynthesis, which contribute to the level of storage oil in oilseed rape (Ruuska et al., 2004) and *Arabidopsis* (Liu et al., 2017) and to the seed weight of soybean (Cho et al., 2023). In some green embryos, photosynthesis has also been linked to the synthesis of ATP and NADPH for the production of complex carbohydrates, fatty acids, and proteins (Asokanthan et al., 1997; Wu et al., 2014). However, other *Arabidopsis* studies using chemical inhibitors of embryonic photosynthesis (Allorent et al., 2015) or inducible photosynthesis-deficient mutants showed that the seed oil and protein content was not changed, but germination and early growth from the seeds was affected (Sela et al., 2020). The effect of pod/seed photosynthesis on seed composition therefore remains unclear.

An objective of this study is to screen for low chlorophyll soybean mutants that have higher levels of seed protein without compromising yield, potentially due to the mutant facilitating the redirection of nitrogen resources towards enhancing seed composition. We assessed the seed composition and yield of twenty-five low chlorophyll mutant varieties sourced from the USDA Soybean Germplasm Collection. One low chlorophyll Lincoln mutant (PI548210) showed a higher concentration of seed protein while maintaining yield compared to its dark green Lincoln parent (PI548362). Additionally, another low chlorophyll *Y11/y11* mutant (PI547555) had a lower oil concentration but comparable yield to its dark green Clark parent (PI548533), and also reduced photosynthesis in the pods and seeds. We hypothesize that reduced chlorophyll levels in pods and seeds may hinder pod photosynthesis, thus affecting seed composition. Our study explores the impact of low chlorophyll mutations on yield, seed composition, and pod/seed photosynthesis, while also exploring correlations between chlorophyll levels in leaves, pods, and seeds.

Hypotheses

1. We hypothesized that light green mutants would reallocate nitrogen from chlorophyll synthesis into protein in the seeds compared to the dark green parent.
2. We hypothesized that pod and seed photosynthesis is lower in the low chlorophyll mutant compared to the dark green parent.
3. Both concepts should result in lower seed oil concentration and higher protein concentration in the mutants compared to their dark green parents.

## RESULTS

### Two low chlorophyll mutants exhibit comparable yield but differences in seed composition compared to their dark green parents

We measured yield (per plant) and seed composition in various low chlorophyll mutant varieties to explore potential correlations with leaf chlorophyll levels. These varieties, primarily natural mutants from the USDA Soybean Germplasm Collection, have varying levels of leaf chlorophyll, and the genetic basis for the phenotype in most cases remains unknown. In our study, a total of 36 soybean varieties were planted, including 25 low chlorophyll mutants and 11 dark green parents (Supplemental Figures 1-4). The low chlorophyll mutants had diverse leaf chlorophyll levels in the 2021 Illinois field, ranging from 11 to 51 in SPAD value (Supplemental Figure 5).

While some low chlorophyll mutants appeared stunted and unhealthy (e.g., PI560909 and PI548265 in Supplemental Figures 1-4), others (e.g., *Y11/y11* and Lincoln mutants in Figure 1 and Supplemental Figures 1-4) were a similar size and growth habit to their dark green parents. We did not observe a strong correlation between leaf chlorophyll levels and seed yield or seed composition; the R^2^ values for yield (0.07), concentration of oil (0.1), and concentration of protein (0.04) were all notably low (Supplemental Figure 5). Additional measurements of amino acids and fatty acids failed to reveal any significant correlation with leaf chlorophyll levels (Supplemental Figure 6).

Recognizing potential genetic background variations in soybean affecting yield and seed composition, we focused on comparing low chlorophyll mutants and their dark green parents within the same genetic background. Among the 25 low chlorophyll mutants screened, 9 exhibited similar yields compared to their dark green parents (Supplemental Figure 7). Interestingly, two mutants with comparable yields displayed distinct seed composition characteristics: the Lincoln mutant (PI548210) had a 51% lower leaf chlorophyll (*p* = 0.001) and 16.7% lower oil concentration (*p* < 0.001), and protein was 5.4% higher compared to its dark green Lincoln parent PI548362 (*p* < 0.009). Cysteine and methionine were also significantly higher in the Lincoln mutant. Another mutant, *Y11/y11* (PI547555), had 62% lower leaf chlorophyll (*p* <0.001), 3% lower oil concentration (*p* = 0.03), and non-significant differences in yield and protein concentrations compared to its dark green Clark parent (PI548533; Table 1).

Overall, out of 11 soybean cultivar families screened in small-scale field trials, two (Lincoln and Clark) were found to have low chlorophyll mutants with differences in seed composition while maintaining comparable yield. Other low chlorophyll mutants trended towards lower yield and/or no notable difference in seed composition from their parent cultivars.

### The impact of low leaf chlorophyll on soybean oil concentration interacts with pod height and row spacing

Previous research has indicated that seeds in the upper canopy exhibit higher protein content and lower oil levels compared to seeds at the lower canopy (Collins & Carter, 1956; Escalante & Wilcox, 1993; Huber et al., 2016). Our 2021 field study supports this by finding that, at the upper canopy (top 4 nodes), seed oil concentration is significantly lower for both the low chlorophyll *Y11/y11* mutant (PI547555) and its dark green Clark parent (PI548533) (Fig. 2A); however, protein levels did not change significantly. At this upper layer, the dark green parent still has significantly more oil than the mutant; however, at the lower layer (with higher oil) the difference between them is no longer significant (Fig. 2A).

Row spacing also affects seed oil concentration in Clark and its low chlorophyll *Y11/y11* mutant. Seed collected from a 2013 field experiment (Slattery et al., 2017) and analyzed for seed composition here showed oil was 2% lower (*p* = 0.006) in the low chlorophyll mutant compared to its dark green parent, when grown at a normal 38 cm row spacing (Figure 2C). However, narrowing the row spacing to 19 cm, resulting in a denser canopy, reduced the difference in oil concentration between the parent and mutant and removed its significance (Fig. 2C), thus suggesting seed oil production is sensitive to canopy density. There were no other changes to other seed components at either row spacing in 2013 (Supplemental Figures 10-11).

These results suggest that *Y11/y11* (PI547555), a low chlorophyll soybean mutant of Clark (PI548533), exhibits a reduced concentration of soybean oil without impacting yield, influenced by canopy structure and pod height within it.

### The Clark low chlorophyll mutant (*Y11/y11*) has a lower seed oil concentration associated with lower gross photosynthesis in pods and seeds compared to the dark green parent

Our previous results showed that *Y11/y11* had reduced seed oil concentration relative to Clark and that this difference varied with canopy position and row spacing (Figure 2). Because pod and seed photosynthesis has been linked to oil production in other species (Ruuska et al., 2004; Liu et al., 2017), we hypothesized that the pod and seed photosynthesis would be lower in the low chlorophyll mutant compared to the dark green parent, which may contribute to the decrease in oil concentration (Table 1 and Figure 2). Under field conditions, we assessed the rates of net photosynthesis and dark respiration (R_dark_) in pods and calculated the rates of gross photosynthesis for both the low chlorophyll mutant *Y11/y11* (PI547555) and the dark green Clark parent (PI548533) (Figure 3). The results, summarized in Table 2, revealed significant differences: the dark green parent exhibited almost two times greater gross photosynthesis both at light saturation (6.1 vs 3.1 μmol CO_2_ m^-2^ s^-1^ at 2000 μmol m^-2^ s^-1^), assuming pod and seeds receive full sunlight, and incident light (4.5 vs 2.6 μmol CO_2_ m^-2^ s^-1^ at 300 μmol m^-2^ s^-1^), assuming pods are situated beneath the canopy, where the estimated incident irradiance on the pods is approximately 260 μmol m^-2^ s^-1^ (Allen et al., 2009). We did not observe significant difference in net photosynthesis at either light saturation or incident light. The rate of respiration in the dark was nearly two times greater in Clark (PI548533) compared to *Y11/y11* (PI547555); however, the rates of gross photosynthesis are slightly higher than respiration, indicating that both Clark and *Y11/y11* were able to recycle most of the CO_2_ lost to respiration.

We investigated the cause of the lower pod and seed photosynthesis in the low chlorophyll mutant. First, we hypothesized that the low chlorophyll mutant (so named due to leaf chlorophyll levels) may have a lower chlorophyll level in pod and seed tissue too. However, we failed to find a significant difference in chlorophyll levels of immature pods and immature seed tissues between *Y11/y11* (PI547555) and the dark green Clark parent (PI548533; Supplemental Figure 12). We next looked at the operating efficiency of photosystem II (F_q_’/F_m_’) in pods of the low chlorophyll mutant and found it to be, surprisingly, 20% higher than that of the dark green parent (0.384 vs. 0.317; Figure 4 A-C), although there was no difference in the operating efficiency of photosystem II in the seeds (Figure 4 D-F). Finally, a simulation of canopy photosynthesis predicted more light is available in the low chlorophyll mutant, which ought to result in higher pod/seed photosynthesis beneath the canopy (Supplemental Figure 13).

Thus, the reason for the lower gross pod and seed photosynthesis in the low chlorophyll *Y11/y11* (Clark mutant) remains unresolved, suggesting an area for further study due to its apparent influence on reducing seed oil concentration.

These results demonstrate that the oil concentration of soybeans is influenced by pod and seed gross photosynthesis. However, the lower gross photosynthesis is not attributed to lower chlorophyll levels in seeds and pods, nor to the operating efficiency of photosystem II or light environments.

### The level of leaf chlorophyll does not correlate with levels of chlorophyll in the pod and seed

We hypothesized that, in mutants known to have lower chlorophyll in the leaves, the levels of chlorophyll in the pod and seed would also be lower. In order to explore the correlation between leaf chlorophyll and pod/seed chlorophyll, we examined chlorophyll levels in these tissues from up to 20 soybean varieties, using either SPAD (leaves) or chlorophyll extraction (pods and seeds). Yet, our investigation did not reveal a strong correlation between leaf chlorophyll and pod/seed chlorophyll at two seed developmental stages, either in the greenhouse (Figure 5) or in the field (Supplemental Figure 14). R^2^ values ranged from 0.088 to 0.28, suggesting low chlorophyll varieties (described based on their most prominent chlorophyll-containing feature, leaves) do not necessarily have proportionately low chlorophyll in their immature pods and seeds compared to dark green varieties.

### Levels of chlorophyll and expression of genes in the chlorophyll synthesis pathway are substantially lower in pods and seeds compared to leaves

The concentrations of chlorophyll in pods and seeds were 5- to more than 100-fold lower than those in leaves (Supplemental Tables 1 and 2), irrespective of variety, indicating that chlorophyll synthesis is substantially less active in pod and seed tissues. To further examine tissue-specific chlorophyll synthesis, we analyzed the expression of 58 genes encoding 16 enzymes involved in the chlorophyll biosynthesis pathway using publicly available soybean transcriptome datasets containing roots, flowers, nodules, young leaves, seven stages of seed development, and three stages of pod development (Severin et al., 2010).

Most genes in the chlorophyll common pathway and Mg-branch exhibited substantially lower expression levels in pods and seeds than in young leaves (Figure 6 and Supplemental Data). Glutamyl-tRNA reductase (GluTR), glutamate 1-semialdehyde aminotransferase (GSA-AT), protoporphyrinogen IX oxidase (PPO), Mg-chelatase (Mgch), and protochlorophyllide reductase (POR) showed up to 30-, 21-, 11-, 53-, and 46-fold lower expression, respectively, in pods and seeds relative to young leaves (Figure 6 and Supplemental Data “Gene expression”). In contrast, genes involved in the heme branch pathway showed a different expression pattern, with several genes exhibiting relatively high expression in pods or root nodules (Figure 6 and Supplemental Data). Two genes, one ALA dehydratase gene (ALAD, Glyma.07G032000) and one coproporphyrinogen III oxidase gene (Glyma.14G003200), exhibited similar or higher expression levels in root nodules than in leaves (Figure 6 and Supplemental Data).

To investigate why leaf chlorophyll levels did not correspond to chlorophyll levels in pod and seed tissues in some mutants (Figure 5), we further examined the expression pattern of ChlI1a (Glyma.13G305600), the Mg-chelatase subunit gene mutated in *Y11/y11* (PI547555) (Campbell et al., 2015). ChlI1a expression was highest in young leaves (106 RPKM), moderate in immature pods (33–41 RPKM), and very low in immature seeds (2–8 RPKM) (Supplemental Data “Gene expression”). These results suggest strong tissue specificity in the expression of chlorophyll biosynthetic genes in dark green varieties, which may also be applicable to low chlorophyll mutants given the consistently lower amount of chlorophyll found in all varieties’ seeds and pods compared to leaves.

## DISCUSSION

Low chlorophyll soybean mutants have been proposed as a strategy to improve nitrogen-use efficiency by reallocating nitrogen resources from chlorophyll biosynthesis to seed protein accumulation. In this study, we screened 25 low chlorophyll soybean mutants and identified one line, a Lincoln mutant (PI548210), that exhibited increased seed protein concentration without an apparent yield penalty relative to its dark green parent, Lincoln (PI548362). This finding is notable because improving soybean seed protein content remains challenging due to the well-established inverse relationship between protein concentration on the one hand, and oil concentration and seed yield on the other (Chung et al., 2003; Chaudhary et al., 2015; Bandillo et al., 2015; Kim et al., 2016; Patil et al., 2017).

The physiological basis for this approach may involve altered nitrogen allocation within leaves. Leaves balance carbon assimilation and nitrogen remobilization, processes that strongly influence seed yield and seed nitrogen accumulation, respectively (Havé et al., 2017). Chlorophyll itself represents a substantial nitrogen investment, accounting for approximately 14% of the leaf nitrogen budget (Evans and Clarke, 2019). Modeling studies have suggested that moderate reductions in chlorophyll content could reduce nitrogen demand while maintaining canopy photosynthesis (Song et al., 2017; Walker et al., 2018). Consistent with this concept, a recent tobacco study demonstrated that reduced chlorophyll levels can improve seed nitrogen content without compromising biomass production (Cho et al., 2024).

Although this study was conducted at a relatively small field scale, the identification of a low chlorophyll Lincoln mutant (PI548210) suggests that moderate reductions in chlorophyll may improve soybean seed composition without negatively affecting yield. Future studies should evaluate the stability of this trait under larger-scale and multi-location field conditions and identify the causal mutation underlying the low chlorophyll phenotype. Unlike *Y11/y11* (PI547555, Clark mutant), which had been studied previously and was selected for additional physiological characterization here, the Lincoln mutant (PI548210) was identified during the mutant screen based on its post-harvest seed composition phenotype. Consequently, further analyses, such as immature chlorophyll levels, were not available for the Lincoln mutant.

The low chlorophyll mutants used in this research were selected based on their physical appearance, and most mutant lines, including the Lincoln mutant (PI548210), have an unknown genetic cause for their low chlorophyll phenotype (Boerma and Specht, 2004). Mutations causing low chlorophyll and their pleiotropic effects sometimes impact the yield and growth habits of the plants (Supplemental Figure 1-4 and 7). For example, mutations that disrupt chlorophyll b can impair photosynthesis and photoprotection (Leverenz et al., 1992; Havaux and Tardy, 1997; Kim et al., 2009; Mao et al., 2023). Conversely, the *Y11/y11* Clark mutant (PI547555) which we also studied here has mutations in subunit I of magnesium chelatase (Campbell et al., 2015), which do not adversely affect growth or yield (Slattery et al., 2017; Sakowska et al., 2018).

The impact of pod/seed photosynthesis on oil production may vary depending on the species and can be influenced by factors such as pod location or light exposure, with the canopy structure playing a significant role. Previous studies have yielded contradictory findings regarding whether reduced pod/seed photosynthesis negatively affects seed protein and oil concentration (Ruuska et al., 2004; Schwender et al., 2004; Hua et al., 2012; Liu et al., 2017) or has no discernible effect (Allorent et al., 2015; Sela et al., 2020). In our prior study, we were unable to definitively conclude the effect of pod/seed photosynthesis on seed oil due to covered pods, resulting in minimal photosynthesis, reducing oil concentration in one variety but not the other (Cho et al., 2023). The present study provides evidence consistent with a role for pod/seed photosynthesis on seed oil by demonstrating that the low chlorophyll *Y11/y11* Clark mutant (PI547555) exhibited a reduced concentration of soybean oil without impacting yield, alongside lower gross photosynthesis (Figure 2-3, Tables 1 and 2). Furthermore, previous studies showed the effect cannot be observed under the denser canopy conditions such as narrow row and lower canopy (Slattery et al., 2017; Figure 2C), suggesting the effect of pod/seed photosynthesis interacts with canopy conditions.

We hypothesized that a low chlorophyll mutant would display lower seed and/or pod photosynthetic activity, thereby leading to a decreased seed oil concentration. However, our observations revealed that the low chlorophyll *Y11/y11* Clark mutant (PI547555) exhibited comparable chlorophyll levels in pod and seed tissues (Supplemental Figure 12) to its dark green Clark parent (PI548533). Moreover, the low chlorophyll mutant did not exhibit lower operating efficiency of PSII (Figure 4) or light availability (Supplemental Figure 13). One possible explanation is that biochemical photosynthetic capacity, such as Rubisco carboxylation capacity (V_cmax_) or the electron transport rate (J_max_), may be lower in the pod and seed tissues of the low chlorophyll mutant, as demonstrated by Walker et al. (2018) in leaves.

Huber et al. (2016) reported increased protein concentration at the expense of oil in seeds throughout the canopy after the removal of neighboring plants; and similarly, in this study we found that low chlorophyll mutants grown in a 2013 study in wider rows, with a less dense canopy, had significantly less oil than the parent, in comparison to a denser canopy where the low chlorophyll mutant had comparable oil to the parent (Slattery et al., 2017; Figure 2). Huber et al. (2016) proposed that increased light energy to elevate temperature at most leaf positions could reduce the level of oil concentration at lower nodes, and indeed other studies have found that the temperature negatively correlates with the oil content in soybean seeds (Wolf et al., 1982; Kane et al., 1997; Naeve and Huerd, 2008). We cannot rule out the possibility that the more abundant light energy under the canopy of low chlorophyll mutants increased the temperature, resulting in less oil accumulation.

The results of this study suggest that tissue-specific regulation of chlorophyll biosynthesis may explain why chlorophyll levels in leaves did not correspond to chlorophyll levels in pods and seeds in several low chlorophyll mutants (Figure 5). Transcriptome analysis showed that most genes in the chlorophyll biosynthesis pathway are expressed at substantially lower levels in pods and seeds than in young leaves in a standard dark green variety (Figure 6). This finding is consistent with the markedly lower chlorophyll concentrations observed in reproductive tissues and suggests that chlorophyll synthesis in pods and seeds is already strongly constrained relative to leaves, irrespective of leaf chlorophyll levels.

This tissue specificity may also explain why the mutation in Mg-chelatase subunit ChlI1a in *Y11/y11* Clark mutant (PI547555) had a pronounced effect on leaf chlorophyll levels but comparatively limited effects in pods and seeds. Because ChlI1a expression is already low in immature seed tissues, chlorophyll synthesis in those tissues may be limited by other steps in the pathway rather than by ChlI1a itself. More broadly, these results suggest that regulation of chlorophyll accumulation differs substantially among soybean tissues and that leaf-specific reductions in chlorophyll do not necessarily predict chlorophyll levels in reproductive tissues.

In summary, our study delved into the effects of low chlorophyll mutations on various aspects of soybean cultivation, including yield, seed composition, and pod and seed photosynthesis. The low chlorophyll Lincoln mutant (PI548210) showed a higher concentration of seed protein without affecting yield compared to its dark green parent (PI548362). On the other hand, the *Y11/y11* Clark mutant (PI547555) demonstrated a lower concentration of seed oil without impacting yield, alongside lower photosynthetic activity (Figures 3-5 and Table 2) and comparable chlorophyll levels (Supplemental Figure 12) in pod and seed tissues. These findings suggest that the oil concentration of the soybean is influenced by pod and seed photosynthesis, which is in turn influenced by pod height and row spacing (Figure 2). Chlorophyll levels in the leaf do not necessarily correlate with those in the pod and seed of low chlorophyll mutants, possibly due to substantially low expression of chlorophyll synthesis genes in these tissues.

## METHODS

### Field conditions

Thirty-six cultivars of soybean *Glycine max* were planted in 34.2-cm rows using standard agronomic practices at the University of Illinois Energy Farm field station (40.11°N, 88.21°W, Urbana, IL, USA) on 27 May 2021. Each row (1 block) consisted of ∼10 plants spaced 3.8 cm (1.5 in) apart. There were 4 blocks for each variety. The blocks were spaced 76.2 cm (30 in) apart. There was a border row of Williams 82 (PI518671).

### Seed yield and composition measurements

From the field, all varieties were harvested at maturity on 4 Oct 2021. Each block for each genotype contained up to 10 plants and seeds were pooled from plants within the same block/genotype. Seeds were obtained from all 4 blocks, then further dried in the oven overnight at 50 ℃, then air-dried for at least 2 weeks before NIR measurement. Total weight and the weight of 100 randomly-selected seeds were obtained from each block; from this, an approximate seed count was calculated. Seeds harvested from the field in 2013 (Slattery et al., 2017) came from 3 blocks and 2 row spacings (7.5 in/19 cm or 15 in/38 cm) per variety (Clark PI548533 and *Y11/y11* PI547555).

Near-infrared (NIR) spectroscopy was performed using the Perten DA7250 (PerkinElmer, Shelton, CT, USA) with the company settings for soybean seeds, non-destructively measuring 28 components including protein, oil, 5 fatty acids, and 18 amino acids. See Supplemental Data File for more details.

### Pod and seed photosynthesis measurements

Photosynthesis on attached pods was measured between 1 September and 15 September 2021 at the UI Energy Farm according to Cho et al. (2023). Pods were measured using a LI-6800 (LICOR Biosciences Inc, Lincoln, NE, USA) fitted with a clear-top chamber (LI-6800-12A) using the 3 cm x 3 cm aperture insert. Two small light sources (LI-6800-02), one on the top and the other on the bottom, were attached to the clear-top chamber to allow illumination on both sides of the pod. Pods were placed flat in the chamber to allow full illumination on both sides of the pod. Pods were photographed and projected pod area was estimated using ImageJ (US National Institutes of Health, Bethesda, MD, USA). The pod was allowed to acclimate in the chamber for at least 15 minutes or until the rate of photosynthesis was stable. Reference CO_2_ was set to 400 μmol mol^-1^, irradiance set to 2500 μmol m^-2^ s^-1^, energy balance set to 30 °C, and relative humidity set to 50%. The chamber was set to an over pressure of 0.1 kPa with a flow rate of 1100 μmol s^-1^. Rates of net photosynthesis were measured across a light-response curve which consisted of the following irradiances for both light sources: 2500, 2000, 1600, 1200, 900, 750, 600, 500, 400, 300, 200, 120, 60, 20, 0, 0, 0, 0 μmol m^-2^ s^-1^ with a minimum wait time between changing intensity of 60 s and a maximum wait time of 90 s and the LI-6800 was set to match before each measurement. Gross photosynthesis was estimated by adding the rate of respiration in the dark to net photosynthesis at each irradiance. Light-response curves were measured on four Clark (PI548533) and four *Y11/y11* (PI547555) pods. Following the photosynthesis measurement the pods were removed from the plant to determine chlorophyll content via extraction.

### Chlorophyll measurement

Separated immature pod and seed samples were frozen in liquid nitrogen for at least 10 minutes and stored in the freezer (−80 °C) until they were lyophilized. Chlorophyll content was determined using 100% ethanol extraction (Ritchie, 2006) and microplate spectrometer (Warren et al., 2008). Seeds of 25-100 mg and 100-200 mg fresh weight, and their pods, from 16 varieties were measured from field-grown plants. For leaves, chlorophyll content was measured by SPAD (Soil Plant Analysis Development; Chlorophyll Meter SPAD-502 Plus; Konica Minolta, Japan) on a sun leaf. Five leaves per block (4 blocks, except PI548215, which has data from only 3 blocks) were measured for each variety. Measurements were taken on 29 Jul 2021 for a maximum of 20 replicates per variety. To compare chlorophyll level between tissues, SPAD readings were converted to chlorophyll content using the function from Walker et al. (2018), and using the molecular weight of chlorophyll a (893.489 g/mol) and SLA (308.6 cm^2^/g DW, Slattery et al., 2017). See Supplemental Data File for the calculations.

### Sample collection and chlorophyll fluorescence for photosystem II operating efficiency

For the dark green Clark parent (PI548533) and its low chlorophyll mutant *Y11/y11* (PI547555), immature pods were collected and opened, and the seeds were removed. The immature seeds were weighed to determine the developmental stage (fresh weight range of 100-200 mg). *F_q_’/F_m_’* was measured by a chlorophyll fluorescence (CF) imager (CF Imager, Technologica, UK). Pod- halves (with seeds removed, outer side facing up) were exposed to 260 µmol m^-2^ s^-1^ (light intensity under the canopy from Allen et al., 2009) for 10 min until stabilized then subjected to a burst of 6152 µmol m^-2^ s^-1^ for 800 ms. Seeds were exposed to 78 µmol m^-2^ s^-1^ (light intensity inside the pod from Allen et al., 2009) for 10 min until stabilized then subjected to a burst of 6152 µmol m^-2^ s^-1^ for 800 ms. After the burst, *F’* and *F_m_’* were automatically measured by the software. The images were manually adjusted and *F_q_’/F_m_’* was calculated. Dark green parent Clark (PI548533) had a total of 26 replicates over 3 blocks for pod-halves and 35 replicates over 3 blocks for whole seeds; *Y11/y11* (PI547555) had a total of 8 replicates over 3 blocks for pod- halves and 11 replicates over 3 blocks for whole seeds.

### Transcriptome data analysis

All raw high-throughput transcriptome (RNA-Seq) data came from Soybase (https://soybase.org/soyseq/) originally published by Severin et al. (2010).

### Statistical analysis

Data were analyzed as randomized complete-block designs with all factors treated as fixed effects, while blocks were considered as random effects (α = 0.05). Models were analyzed using mixed effects models with restricted maximum likelihood (REML) and Satterthwaite’s estimate for degrees of freedom (R statistical software, version 4.0.5; lmer function from lme4 package, version 1.1-27.1; least square means differences using the difflsmeans function from lmerTest package, version 3.1-3). P-values were adjusted for multiple comparisons using Tukey’s adjustment, or where appropriate Dunnett’s comparison to control was used, with wild-type as the control.

## Data Availability

All study data are included in the article and/or SI Appendix.

## Acknowledgements

We thank D. Drag, B. Harbaugh, B. Thompson, R. Edquilang, R. Metallo and M. Flack for plant care and management in the greenhouse and field studies; and Y. Ren, D. Nguyen for assistance during laboratory and field work. We thank the staff at the National Soybean Germplasm Collection in Urbana, IL, for curation and distribution of soybean mutants used in this study. We thank J. McGrath and M. Altschuler for critical review of the manuscript. Funding: This work is supported by the research project Realizing Increased Photosynthetic Efficiency (RIPE) that is funded by the Bill & Melinda Gates Foundation, Foundation for Food and Agriculture Research, and U.K. Foreign, Commonwealth & Development Office under grant no. OPP1172157. This work is licensed under a Creative Commons Attribution 4.0 International (CC BY 4.0) license, which permits unrestricted use, distribution, and reproduction in any medium, provided the original work is properly cited. To view a copy of this license, visit https://creativecommons.org/licenses/by/4.0/. This license does not apply to figures/photos/artwork or other content included in the article that is credited to a third party; obtain authorization from the rights holder before using such material.

## Author Contributions

Y.B.C., S.S.S., S.I.J., D.R.O. designed research; Y.B.C., S.S.S., S.I.J. performed experiments; Y.B.C., S.S.S., S.I.J., Y.W., E.A. analyzed data; Y.B.C., S.S.S., S.I.J., Y.W., D.R.O. wrote the paper.

## Reviewers

Interested in soybean seed composition, canopy, chlorophyll

## Abbreviations

Ascension number PI548533 Clark dark green parent

Ascension number PI547555 Y11/y11 Clark mutant

Ascension number PI548362 Lincoln dark green parent

Ascension number PI548210 Lincoln mutant

