## Supplemental Figures and Tables for "Impact of Reduced Chlorophyll Levels in Leaves on Soybean Yield, Seed Composition, Pod/Seed Photosynthesis, and Chlorophyll Levels in Pod and Seed Tissues"

Border Williams

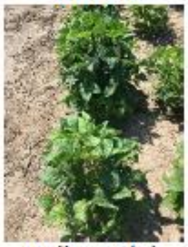

Williams (P) 518671

Richland (M) 548206

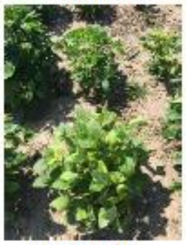

Clark (M) 548217

T239 (M) 548230

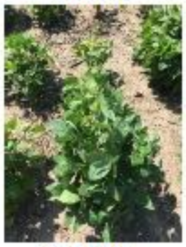

Illini (M) 548211

Peking (P) 548402

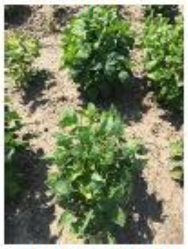

Richland (M) 548207

Peking (M) 548205

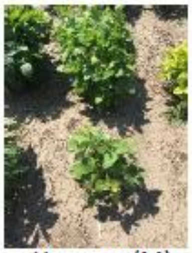

Harosov (M) 547688

Lincoln(M) 548210

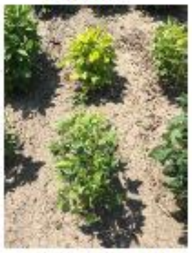

Mandarin (M) 548185

T322 (M) 560910

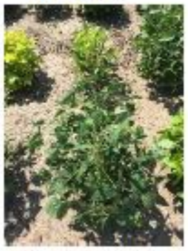

Mandarin (P) 548378

T322 (P) 540553

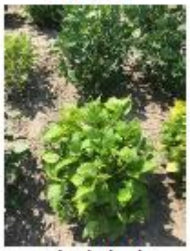

Clark (M) 547555

T239 (P) 548221

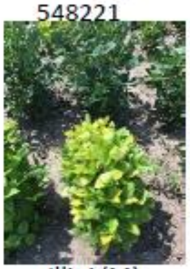

Illini (M) 548172

T322(M) 560911

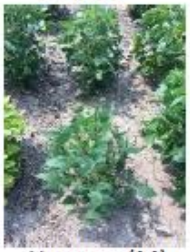

Harosoy (M) 547705

Clark (M) 547418

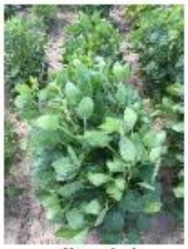

Illini (P) 548348

Richland (P) 548406

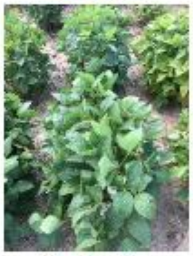

Clark (P) 548533

Clark (M) 547452

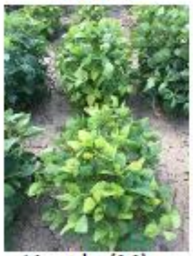

Lincoln (M) 548204

Lincoln (P) 548362

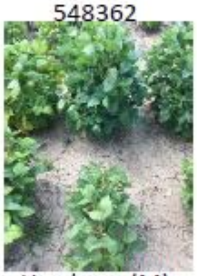

Hawkeye (M) 548216

Hawkeye (P) 548577

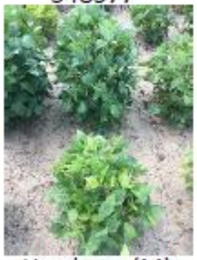

Hawkeye (M) 548215

Harosoy (M) 548229

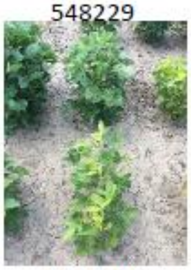

T322 (M) 560909

Williams (M) 548289

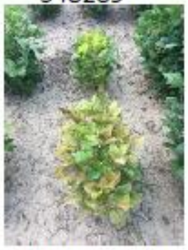

Williams (M) 548265

Williams (P) 548631

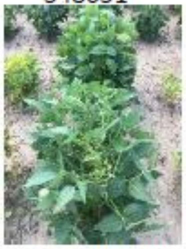

Harosoy (P) 548573

Clark (M) 548231

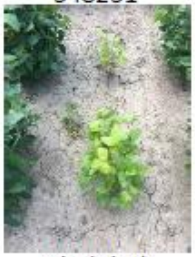

Clark (M) 548260

Border Williams

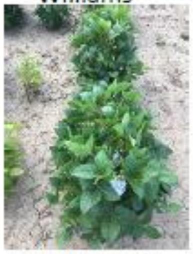

Williams 82 518671

**Supplemental Figure 1. Pictures of low chlorophyll soybean mutants and their parents in the 2021 Illinois field (Block 1).** Each image generally shows two different varieties (labeled top and bottom), clusters of approximately 10 plants.

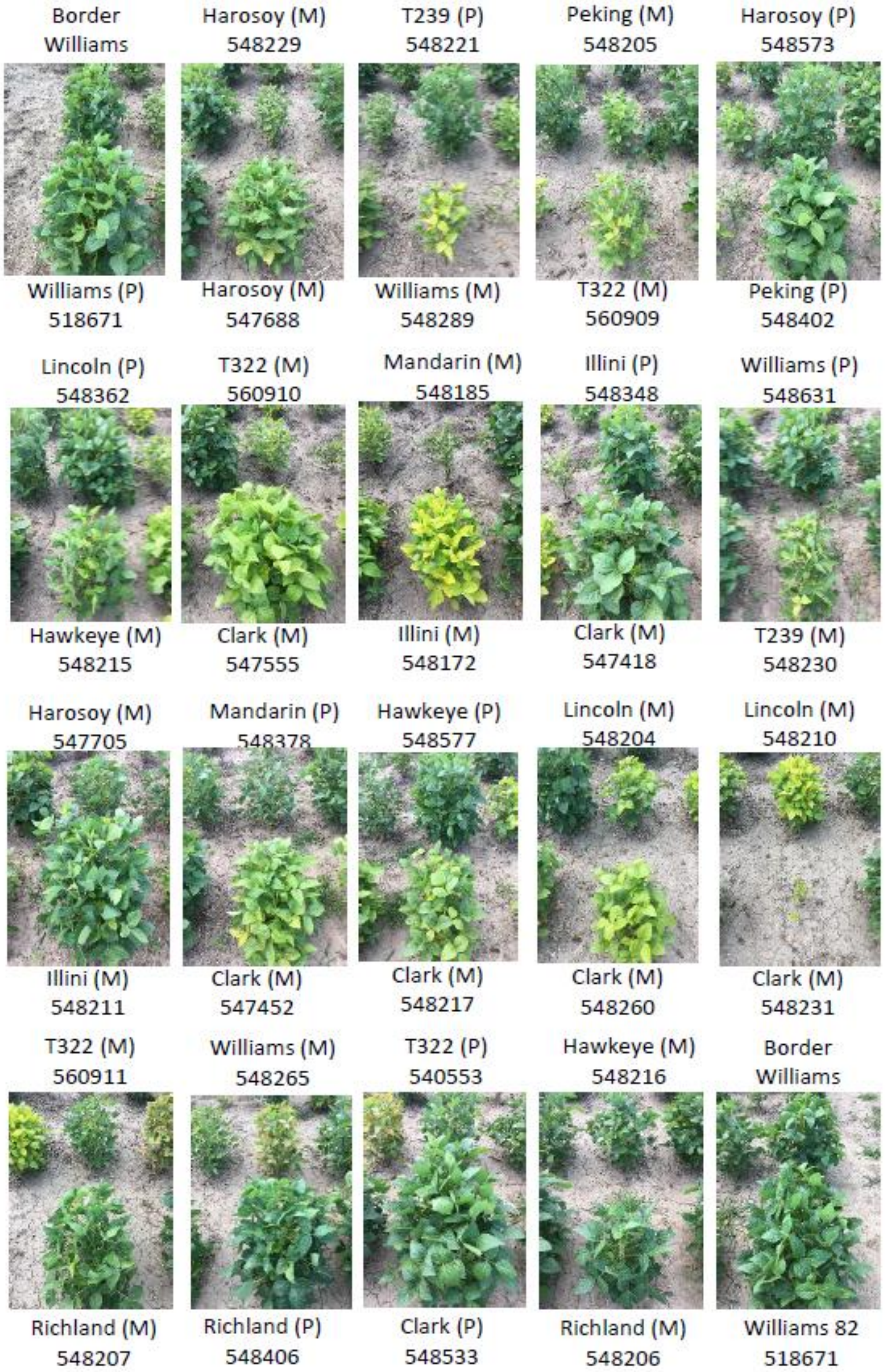


**Supplemental Figure 2. Pictures of low chlorophyll soybean mutants and their parents in the 2021 Illinois field (Block 2).** Each image generally shows two different varieties (labeled top and bottom), clusters of approximately 10 plants.

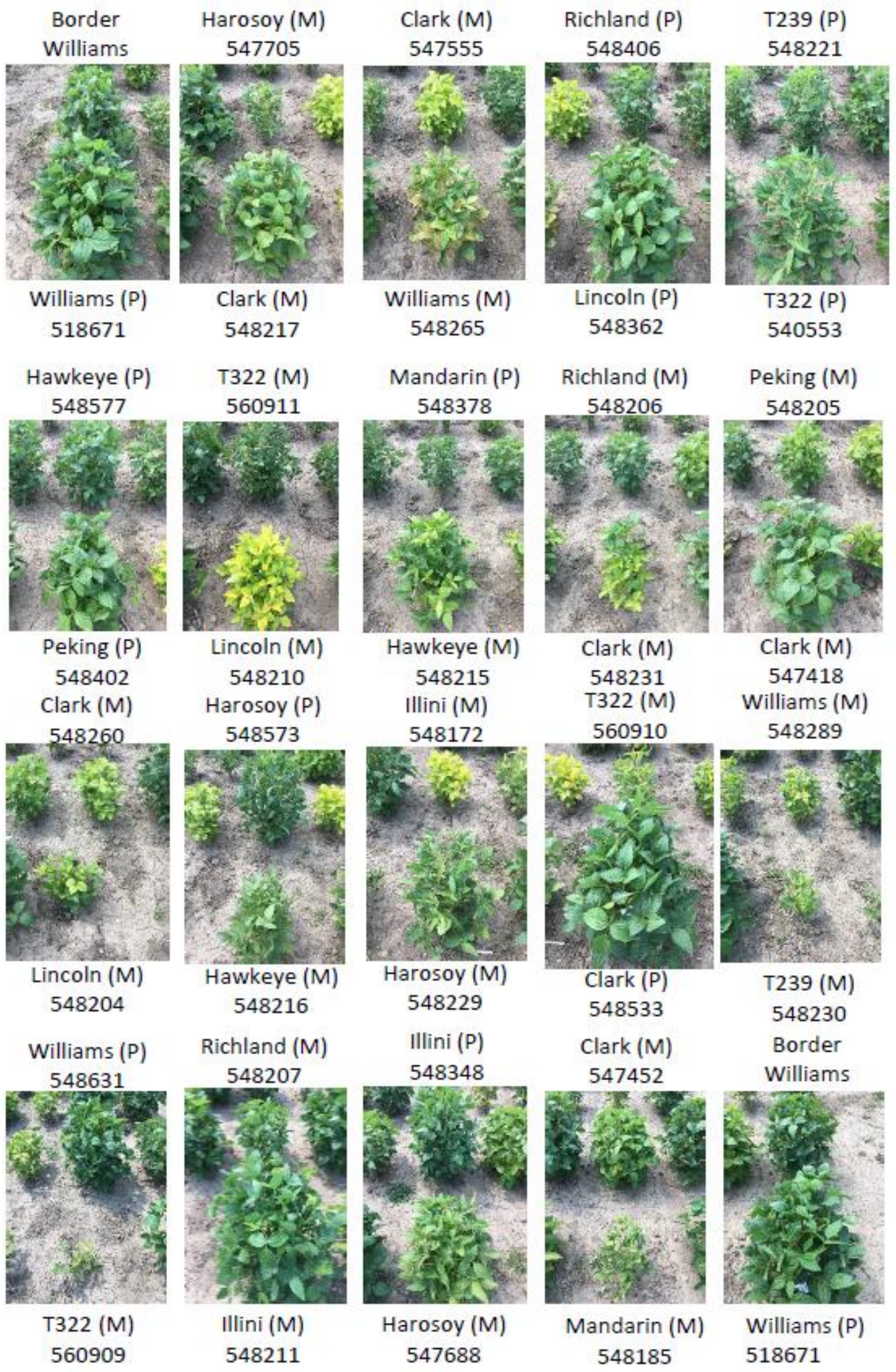


**Supplemental Figure 3. Pictures of low chlorophyll soybean mutants and their parents in the 2021 Illinois field (Block 3).** Each image generally shows two different varieties (labeled top and bottom), clusters of approximately 10 plants.

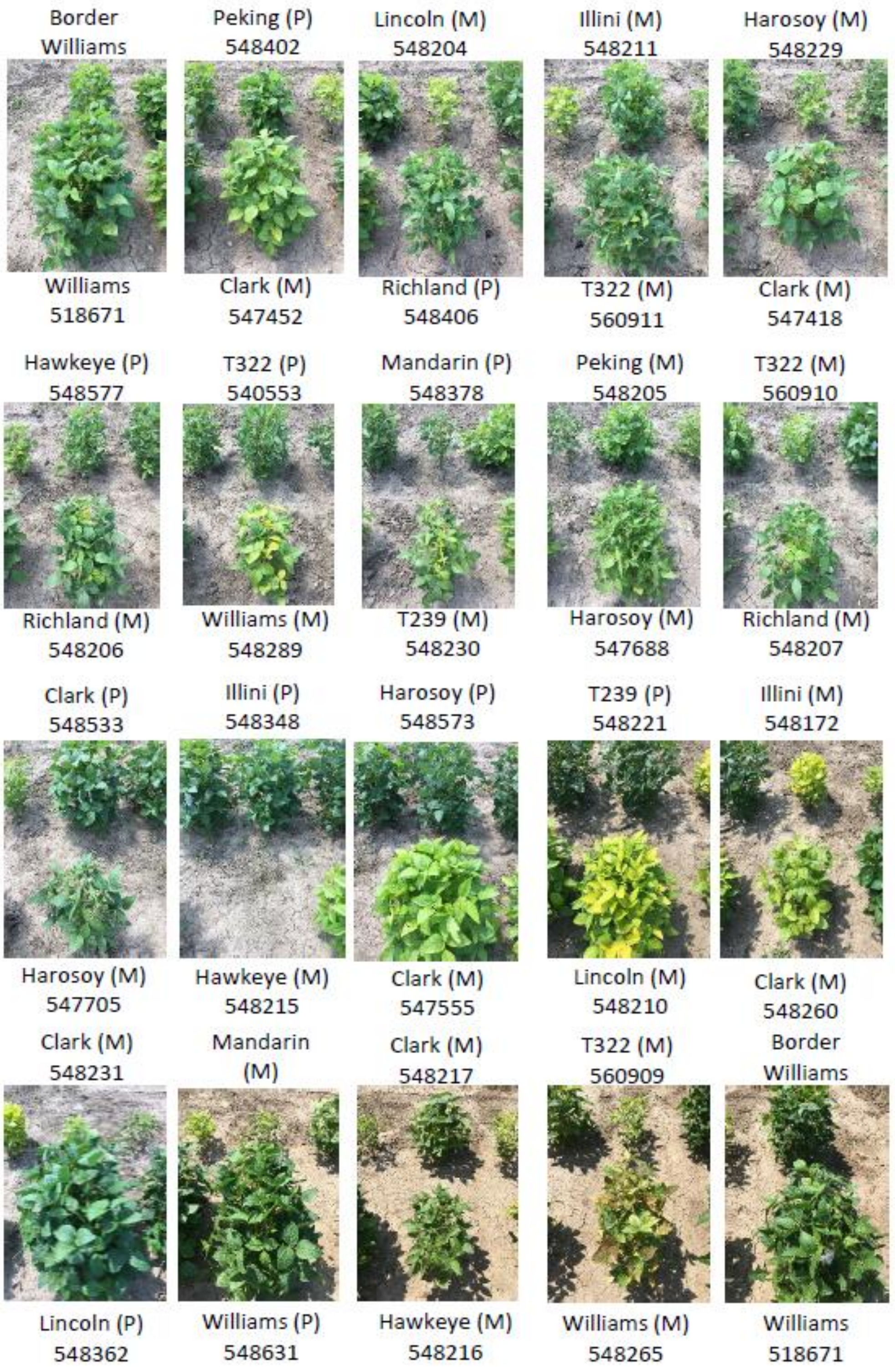


**Supplemental Figure 4. Pictures of low chlorophyll soybean mutants and their parents in the 2021 Illinois field (Block 4).** Each image generally shows two different varieties (labeled top and bottom), clusters of approximately 10 plants.

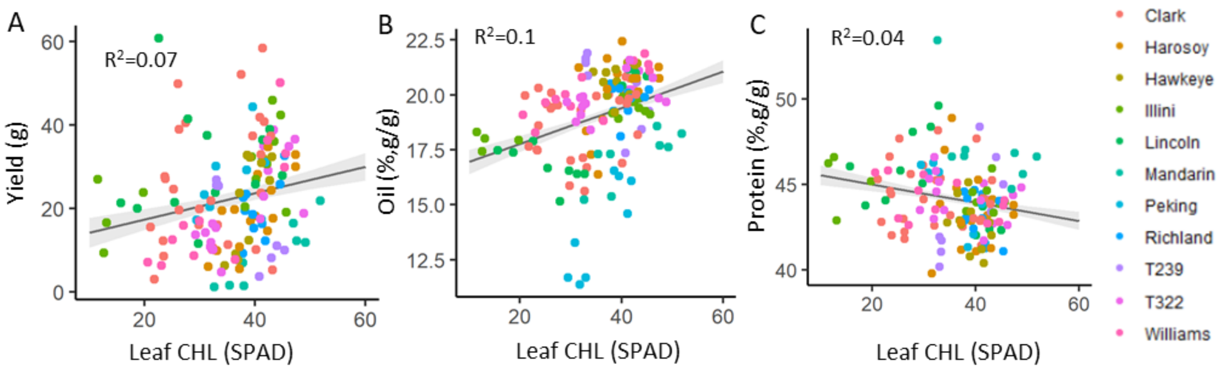


**Supplemental Figure 5. Correlation between the level of leaf chlorophyll and seed yield and major seed components.**

Correlation between the level of leaf chlorophyll (x-axis: SPAD reading) and yield (A), and concentrations of oil (B) and protein (C). Different families (genetic backgrounds) are labelled by different colored dots. Line represents the linear regression model. 25 mutants and 11 parents (36 varieties) and their dark green parents are represented here. R-squared is a coefficient of determination, the percentage of the response variable variation that is explained by the linear model.

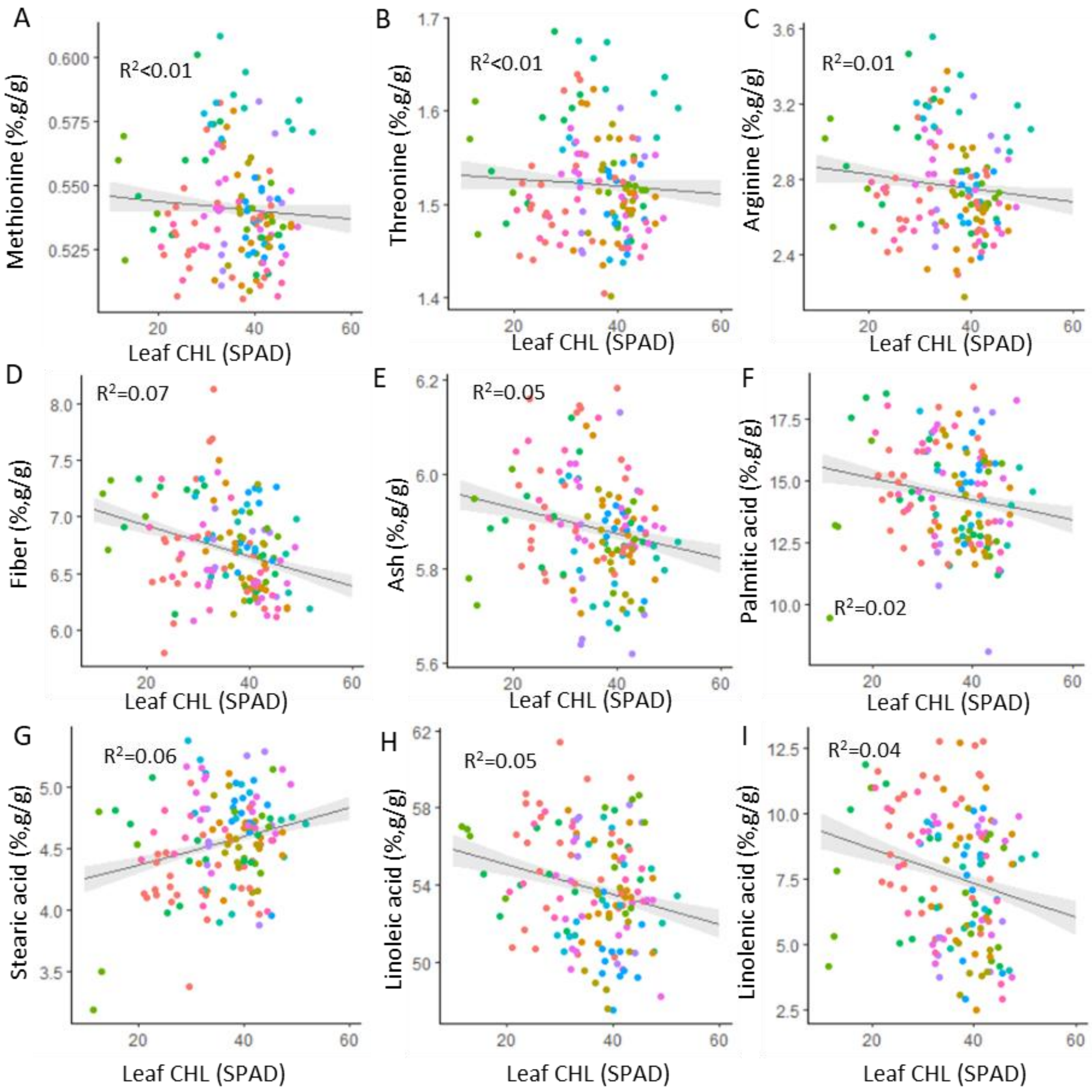


**Supplemental Figure 6. Correlation between the level of leaf chlorophyll (x-axis: SPAD reading) and seed composition.** Different families (genetic backgrounds) are labelled by different colored dots. See Figure 1 for key. Line represents the linear regression model. R-squared is a coefficient of determination, the percentage of the response variable variation that is explained by the linear model.

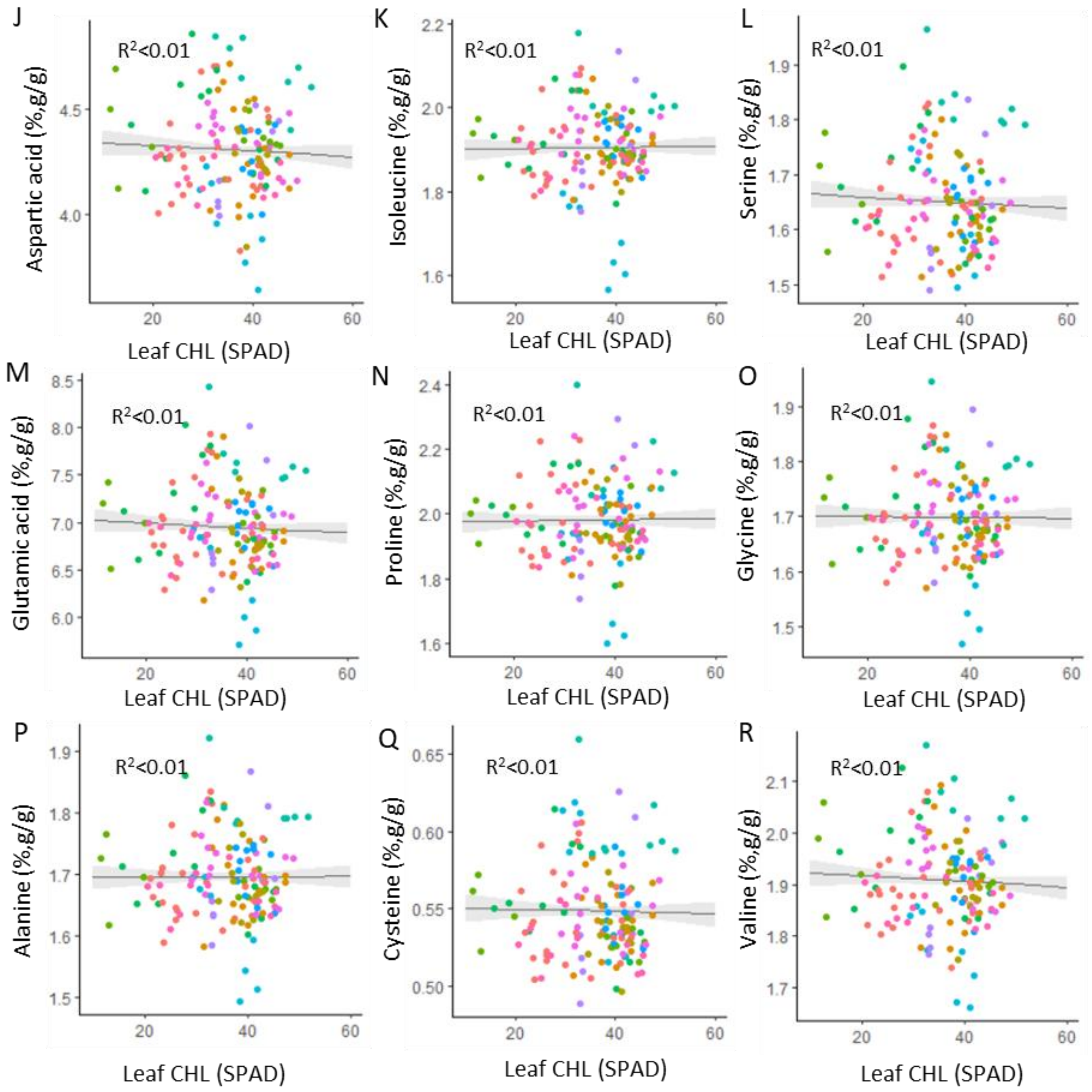


**Supplemental Figure 6 (continued).**

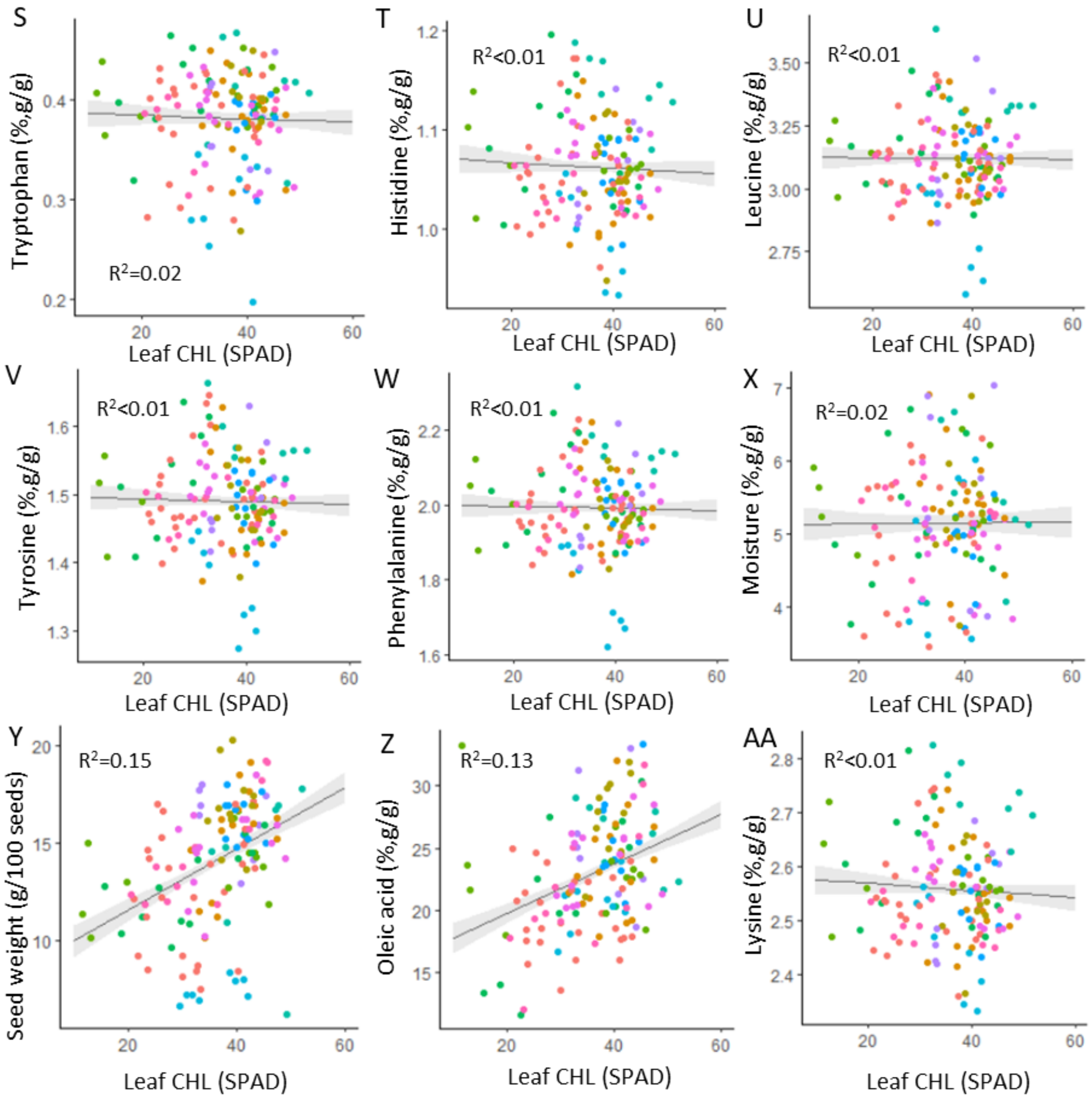


**Supplemental Figure 6 (continued).**

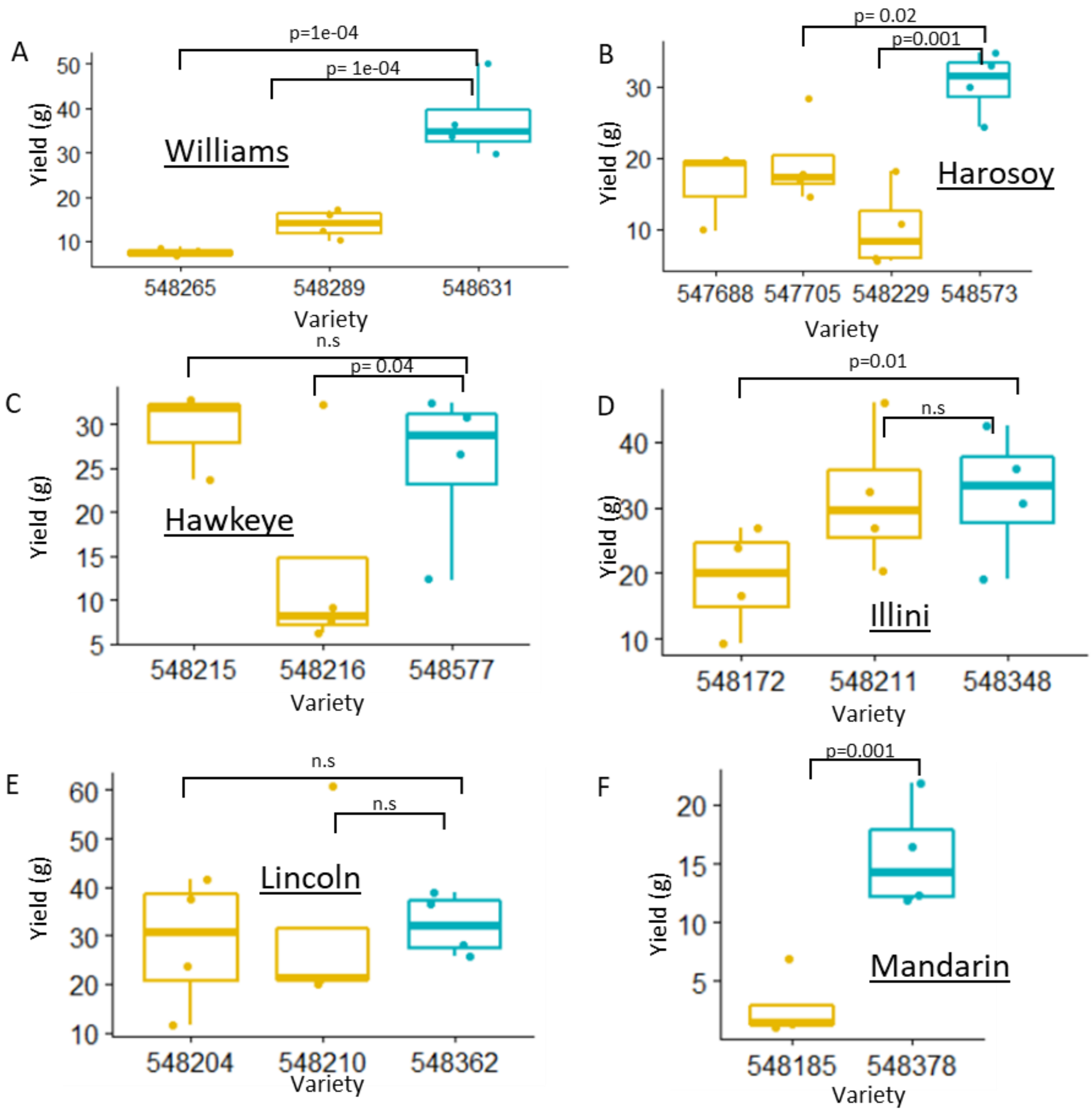


**Supplemental Figure 7. Comparison of yield between low chlorophyll mutants and their parents.**

A. Williams background. B. Harosoy C. Hawkeye. D. Illini. E. Lincoln. F. Mandarin. G. Peking background. H. Richland (548207, N=3 blocks) I. T239 J. T322 K. Clark.

The box plots show the median (central line), the lower and upper quartiles (box) and the minimum and maximum values (whiskers). The statistical analysis was done using ANOVA with linear mixed model (n=4 blocks, alpha=0.05). N.s., non-significant in the analysis. Parent line shown in blue, low chlorophyll mutants in yellow. Yield is average grams of seeds per plant.

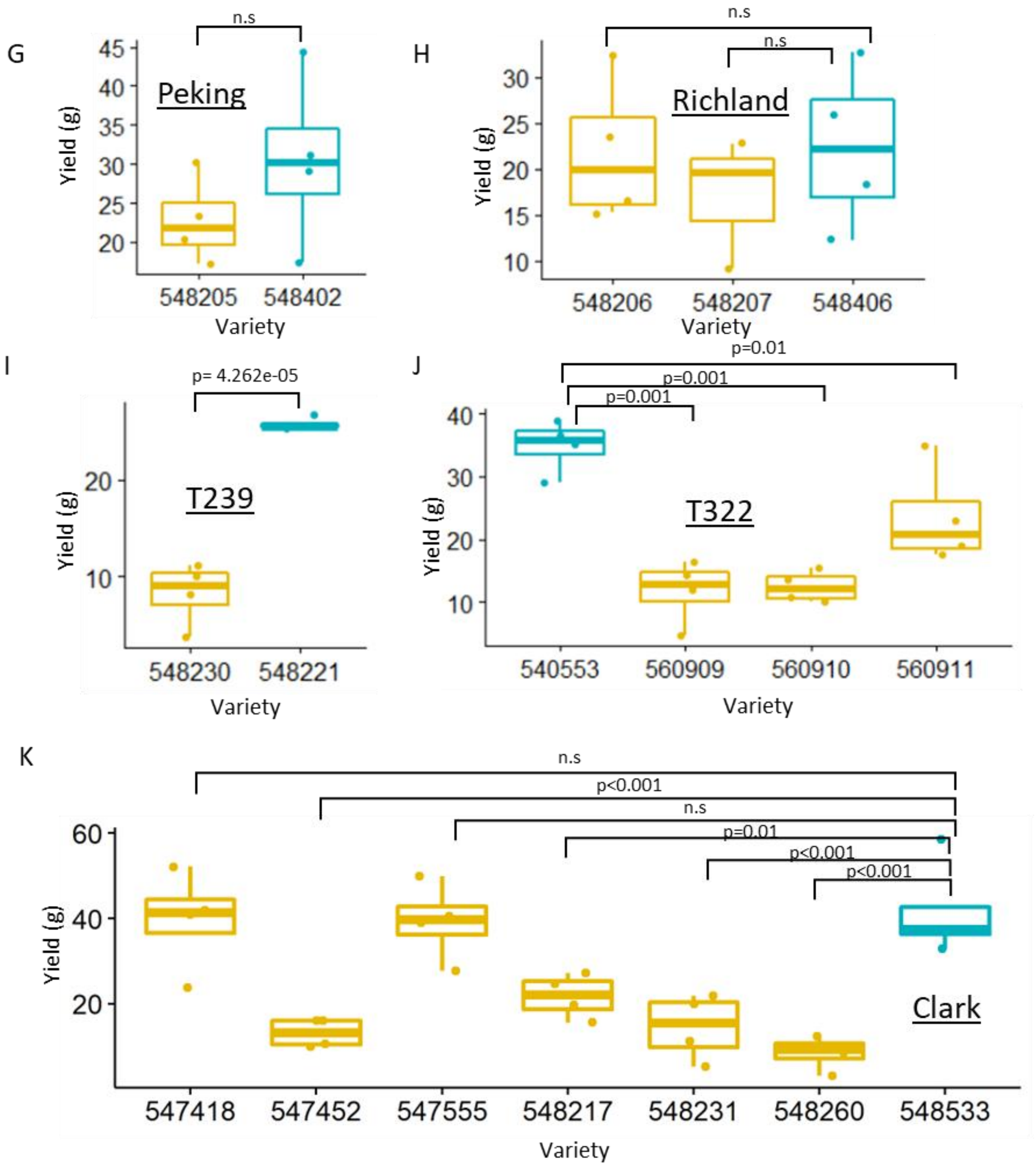


**Supplemental Figure 7 (continued).**

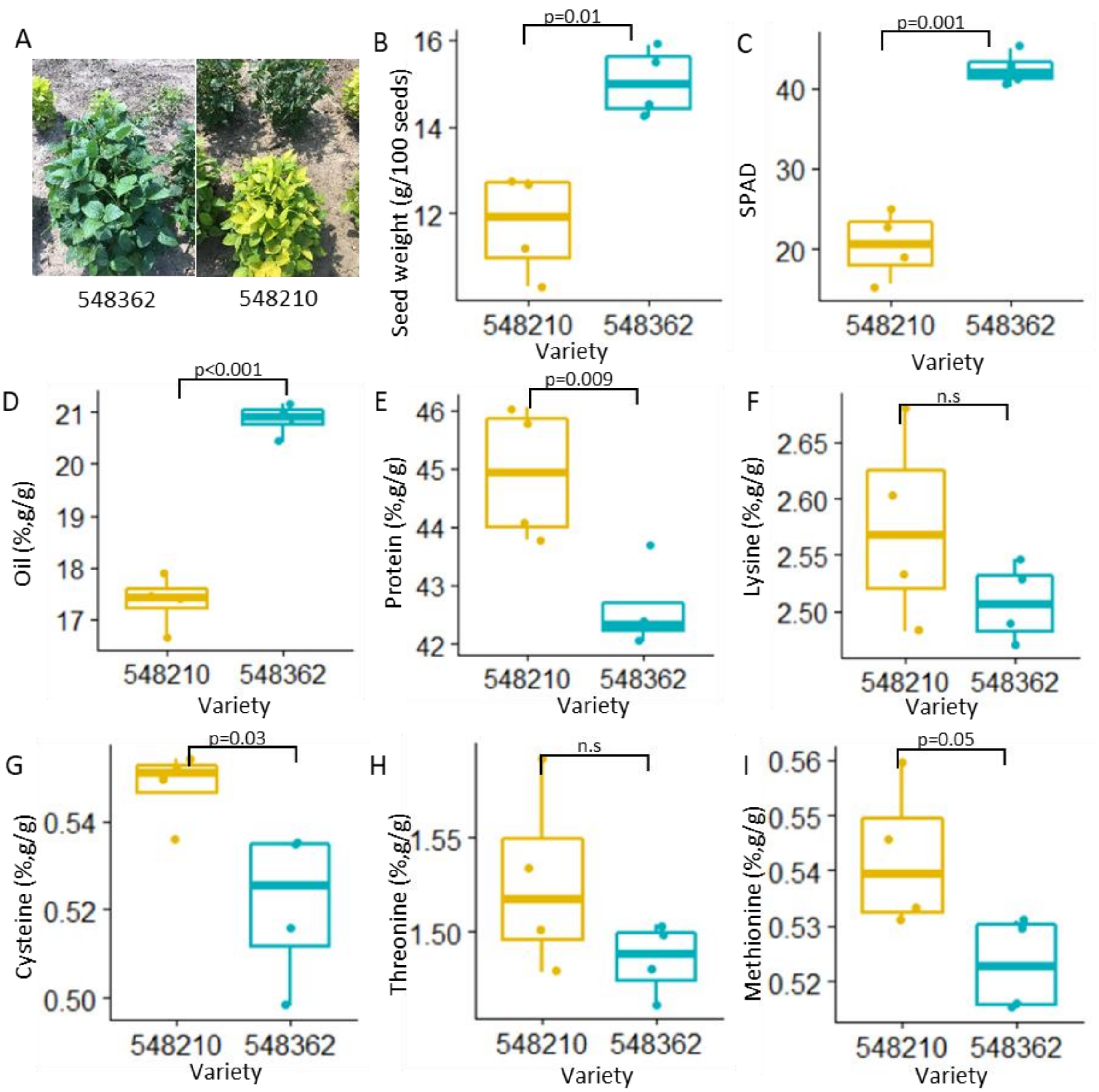


**Supplemental Figure 8. Comparison of yield between low chlorophyll mutants and their parent (Lincoln).** The box plots show the median (central line), the lower and upper quartiles (box) and the minimum and maximum values (whiskers). The statistical analysis was done using ANOVA with linear mixed model (n=4 blocks, alpha=0.05). N.s., non-significant in the analysis. Picture of low chlorophyll mutant (PI548204) and its parent (Lincoln, PI548362) at the 2021 Illinois field (A). Seed weight (B), leaf chlorophyll (C), concentrations of oil (D), protein (E), lysine (F), cysteine (G), threonine (H), and methionine (I).

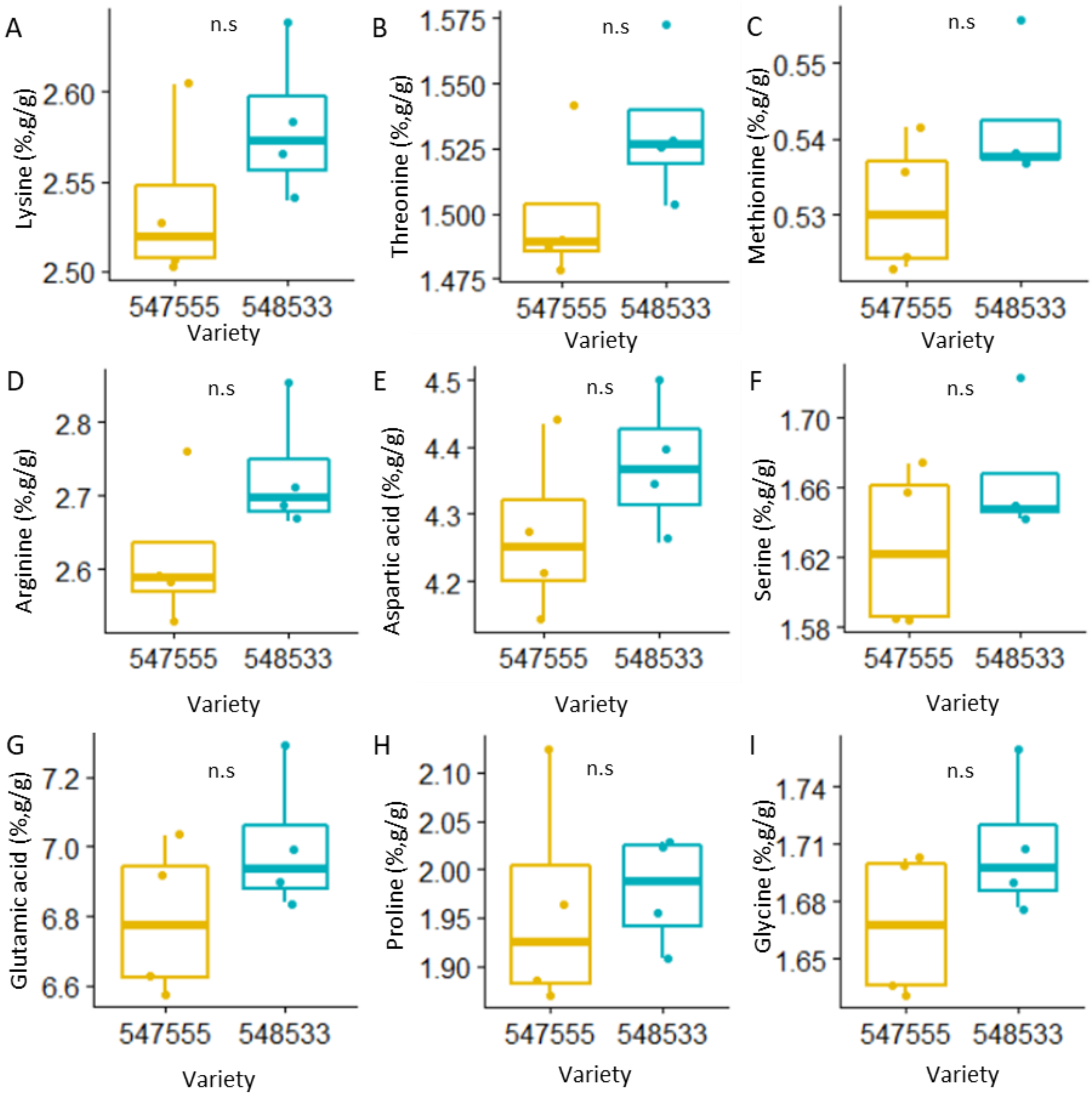


**Supplemental Figure 9. Comparison of seed composition between low chlorophyll mutant (Y11/y11, PI547555) and its parent (Clark, PI548533) from 2021 field.**

The box plots show the median (central line), the lower and upper quartiles (box) and the minimum and maximum values (whiskers). The statistical analysis was done using ANOVA with linear mixed model (n=4 blocks, alpha=0.05). N.s., non-significant in the analysis.

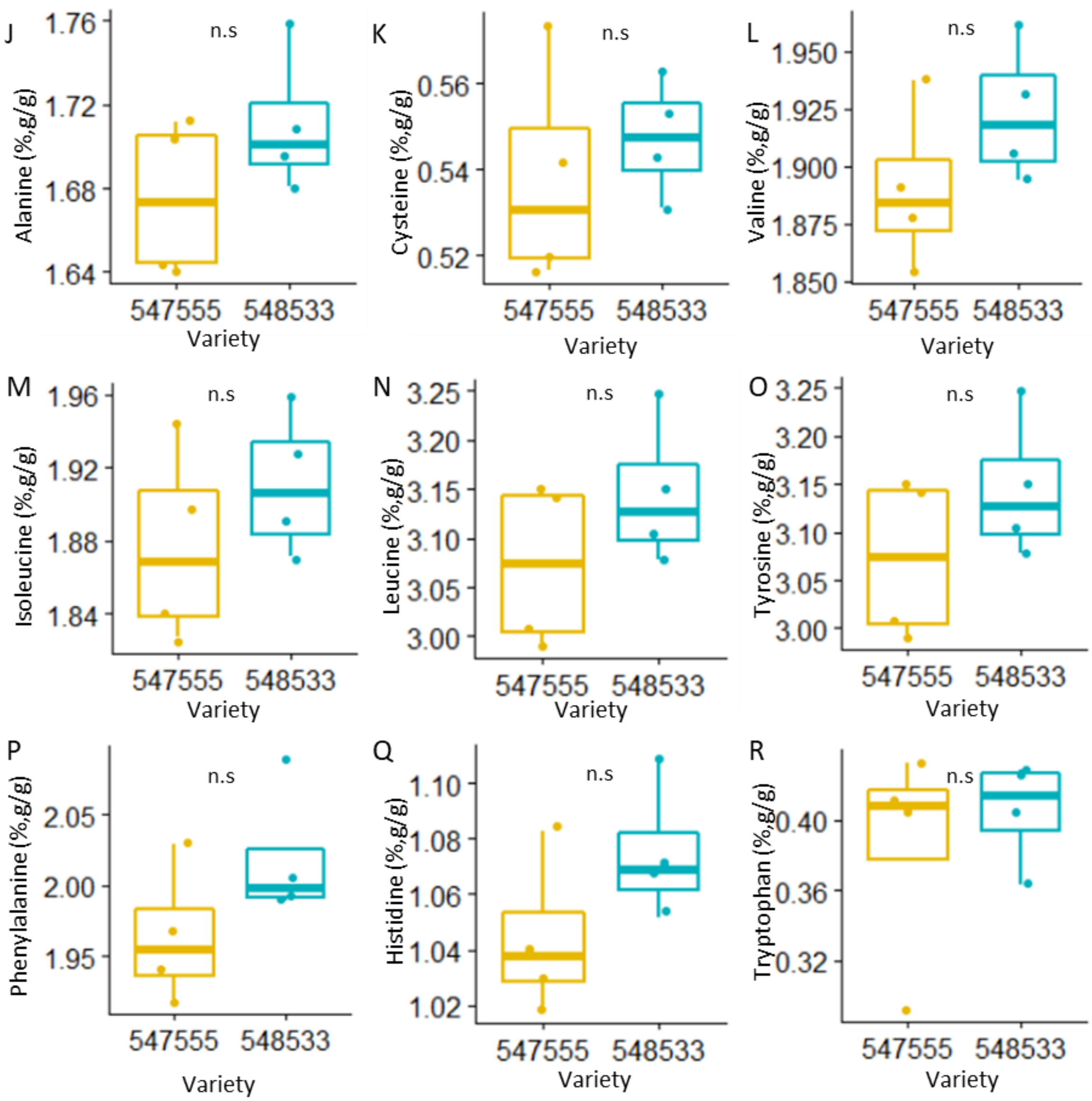


**Supplemental Figure 9 (continued).**

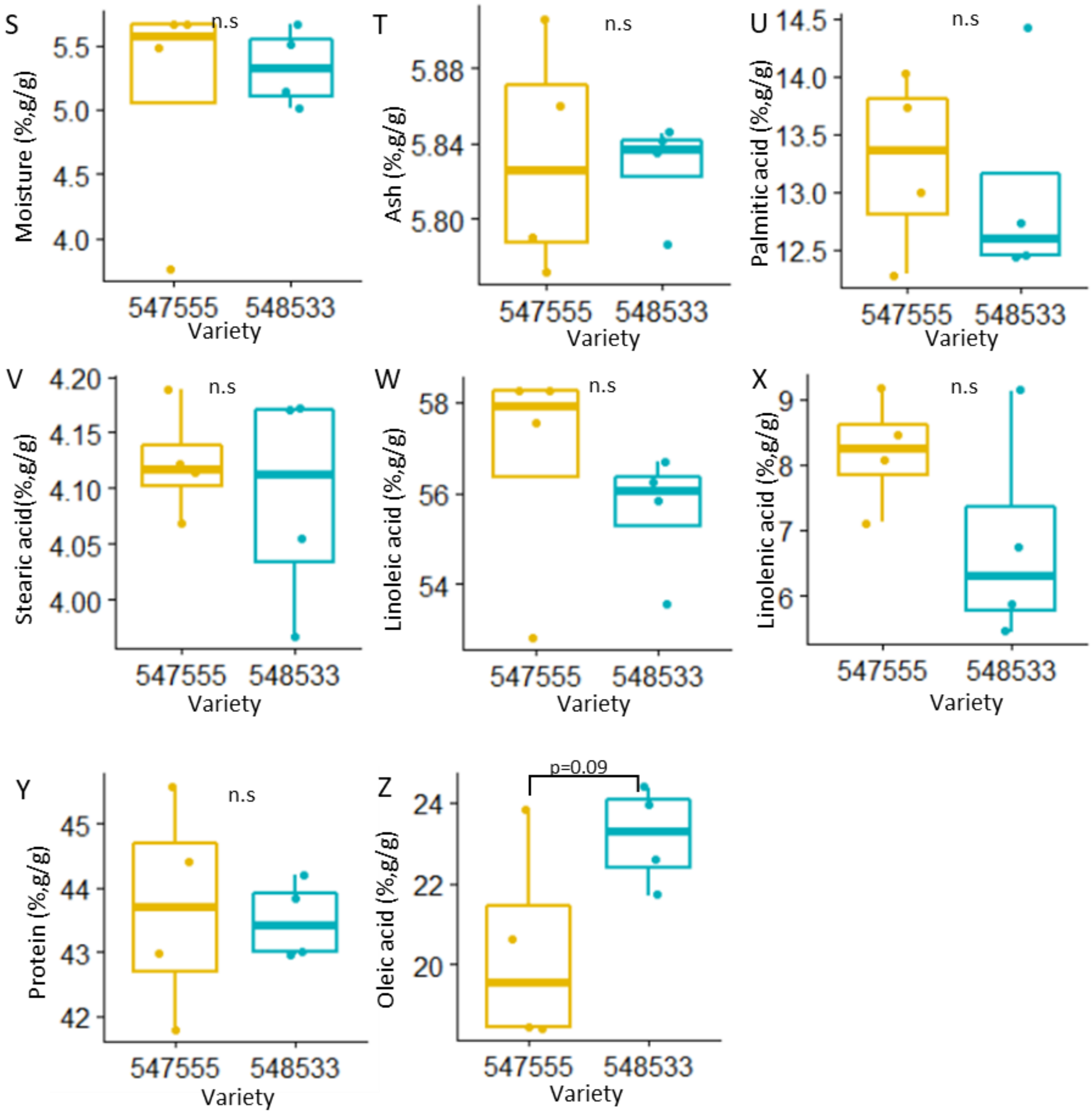


**Supplemental Figure 9 (continued).**

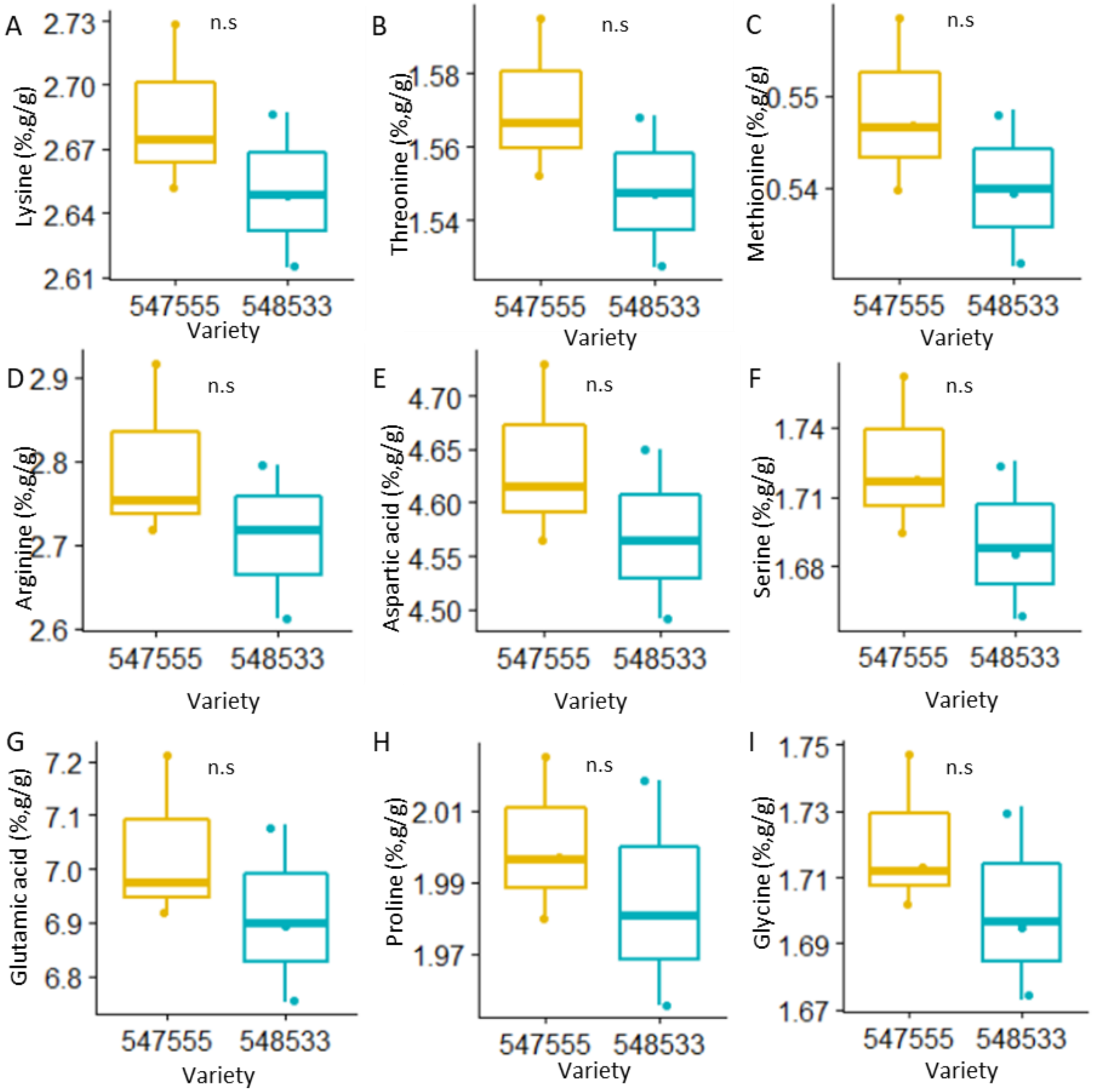


**Supplemental Figure 10. Comparison of seed composition between low chlorophyll mutant (Y11/y11, PI547555) and its parent (Clark, PI548533) from 2013 field (7.5" spacing).** The box plots show the median (central line), the lower and upper quartiles (box) and the minimum and maximum values (whiskers). The statistical analysis was done using ANOVA with linear mixed model (n=3 blocks, alpha=0.05). N.s., non-significant in the analysis.

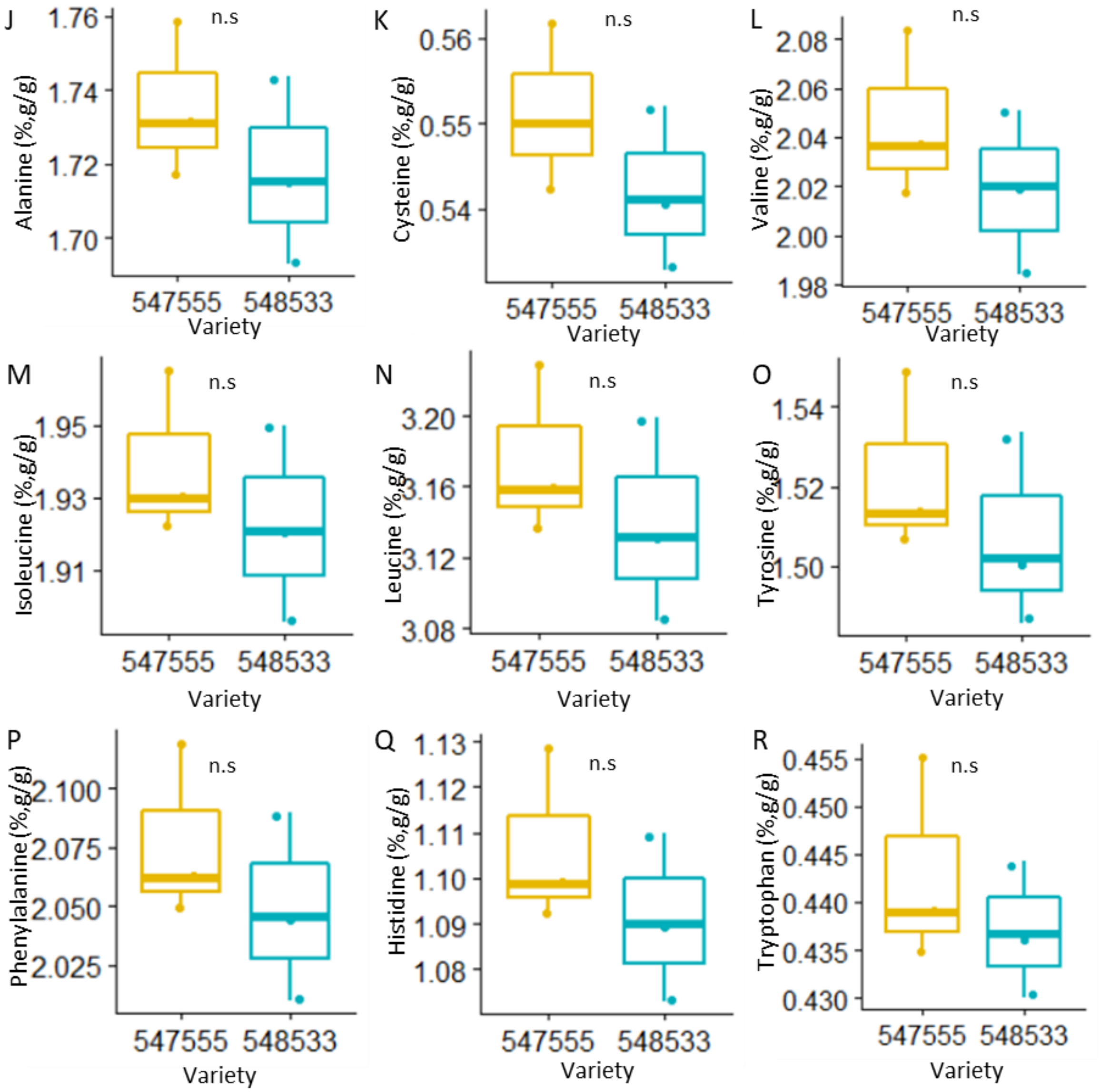


**Supplemental Figure 10 (continued).**

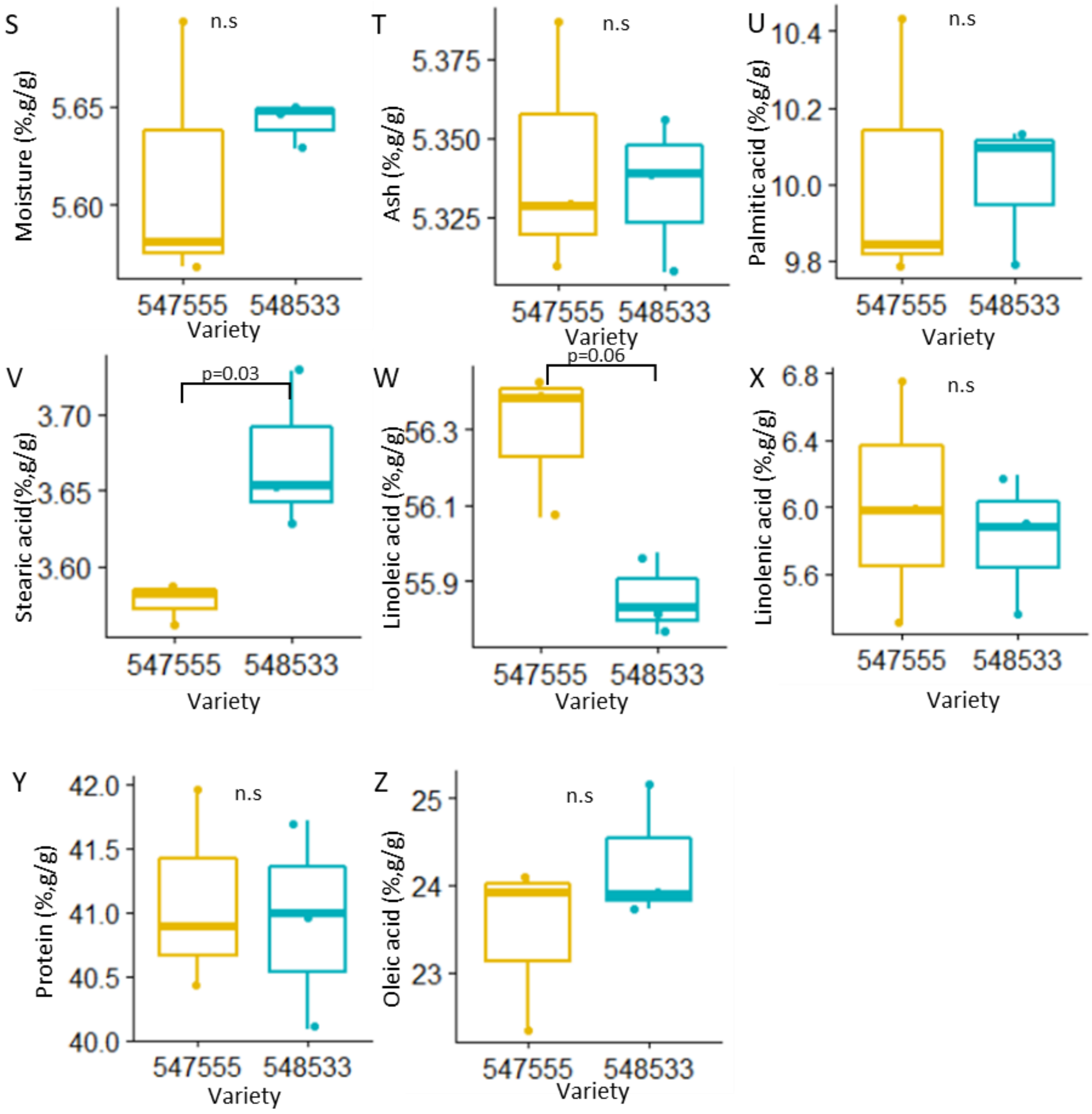


**Supplemental Figure 10 (continued).**

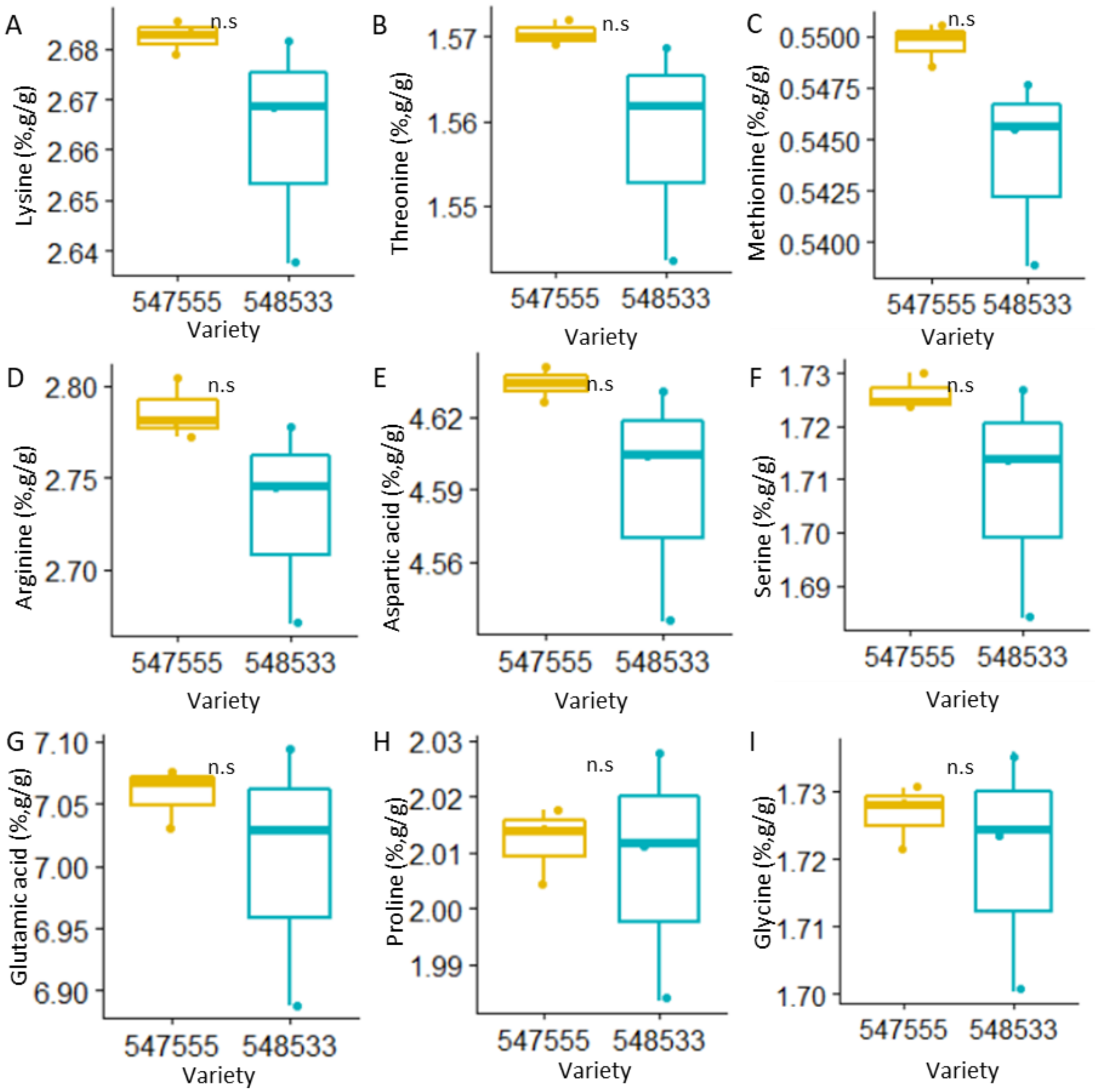


**Supplemental Figure 11. Comparison of seed composition between low chlorophyll mutant (Y11/y11, PI547555) and its parent (Clark, PI548533) from 2013 field (15” spacing).** The box plots show the median (central line), the lower and upper quartiles (box) and the minimum and maximum values (whiskers). The statistical analysis was done using ANOVA with linear mixed model (n=3 blocks, alpha=0.05). N.s., non-significant in the analysis.

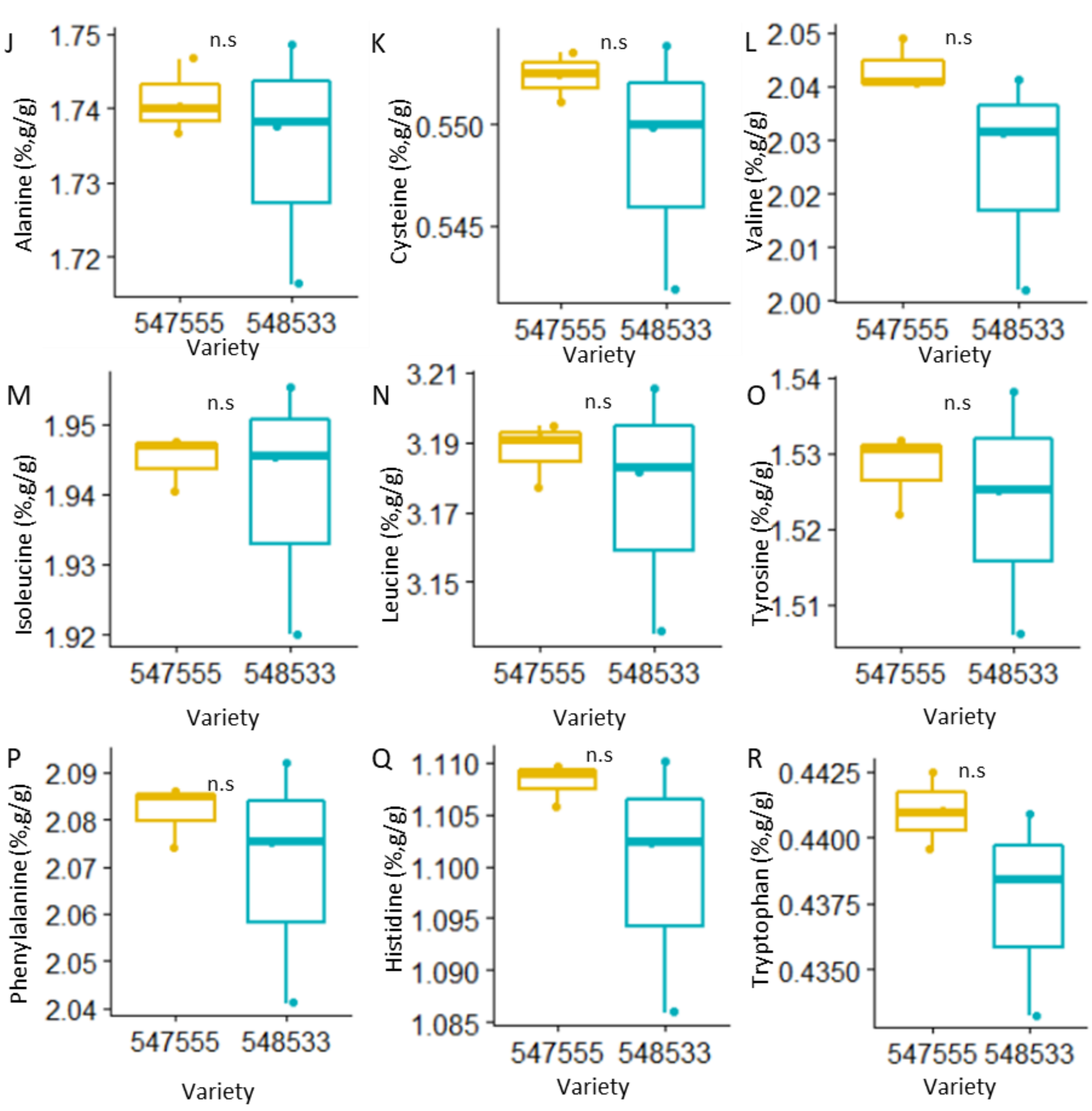


**Supplemental Figure 11 (continued).**

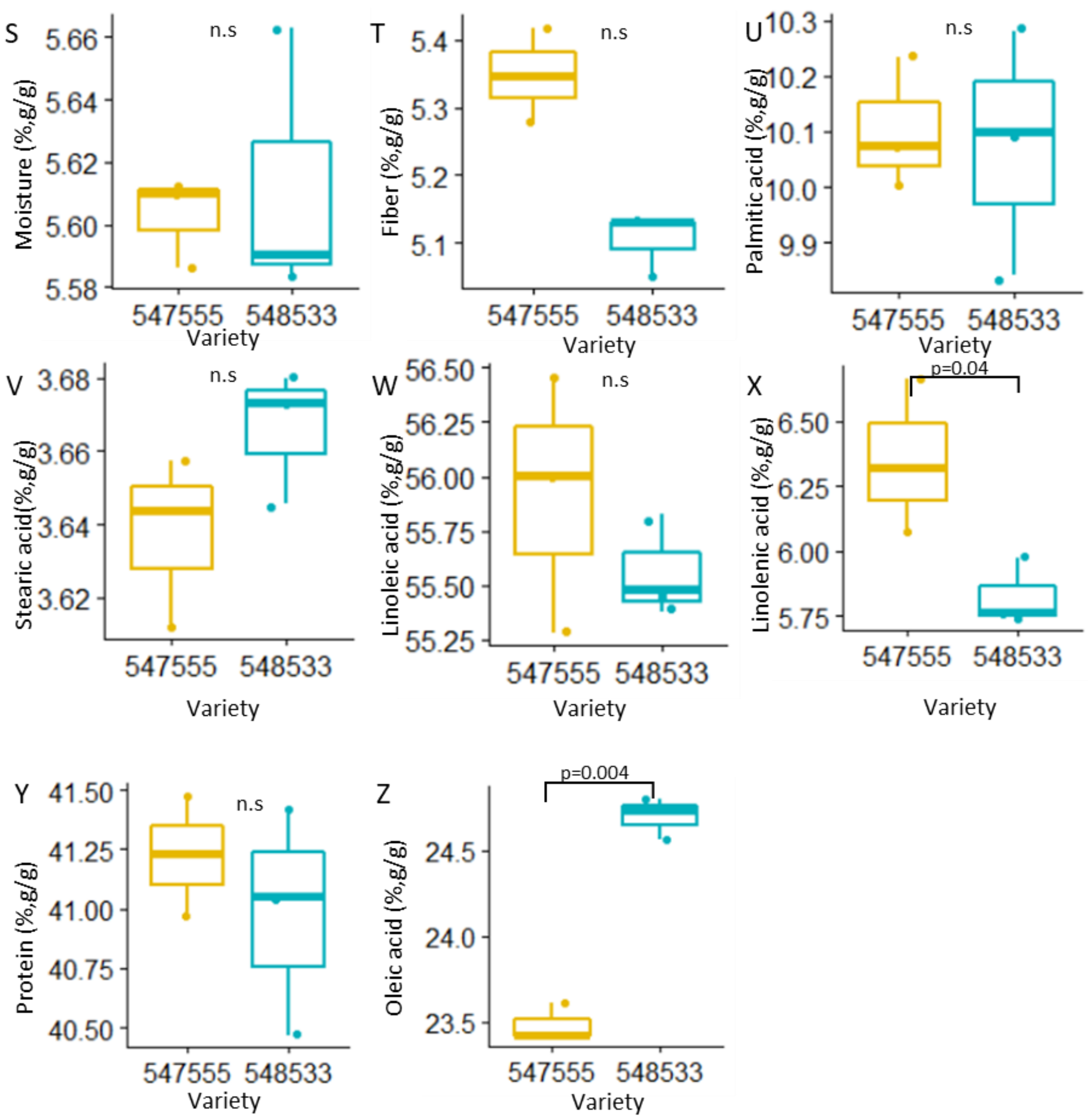


**Supplemental Figure 11 (continued).**

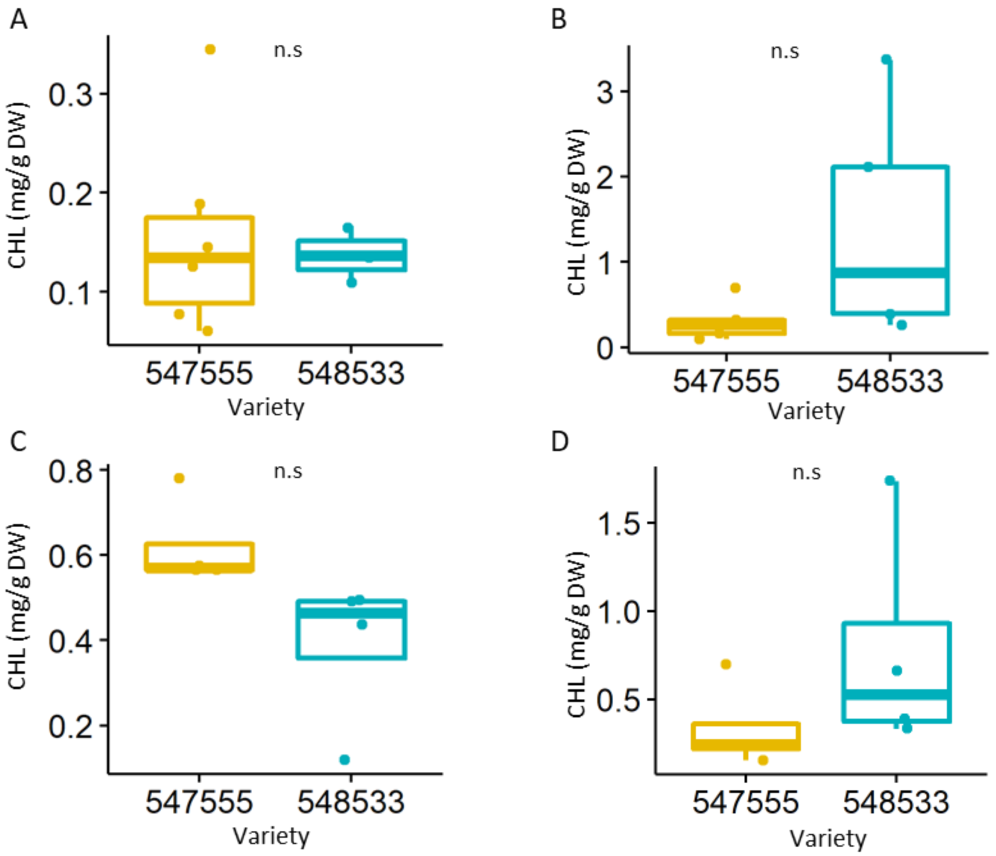


**Supplemental Figure 12. Chlorophyll level of pods and seeds in low chlorophyll mutant (Y11/y11, PI547555) and its parent (Clark, PI548533).** The box plots show the median (central line), the lower and upper quartiles (box) and the minimum and maximum values (whiskers). The statistical analysis was done using ANOVA (alpha=0.05). A-B Field. C-D Greenhouse. N.s, non-significant in the analysis.

A. Chlorophyll level of pods (25-100mg seeds, n=3-6)

B. Chlorophyll level of whole seeds (25-100mg, n=5).

C. Chlorophyll level of pods (100-200mg, n=4).

D. Chlorophyll level of whole seeds (100-200mg, n=4).

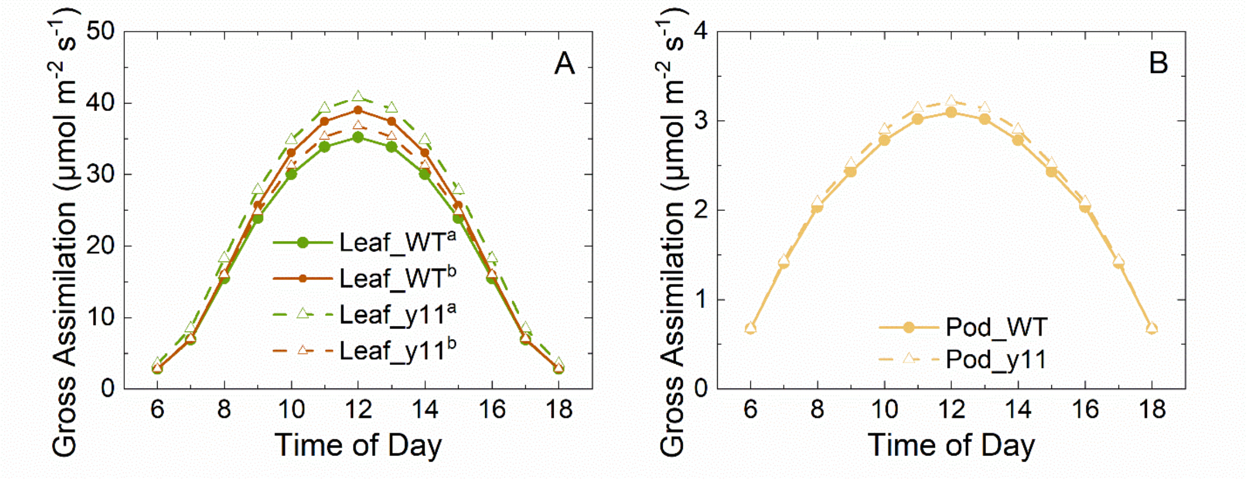


**Supplemental Figure 13. Predicted diurnal gross CO2 assimilation of leaf and pod in the canopy of low chlorophyll mutant (*Y11/y11*, PI547555) and its dark green WT parent (Clark, PI548533) on August 17, 2022.** Model of leaf and pod contribution to assimilation.

A. Gross $CO_2$ assimilation of leaf; **[a]** assumes low chlorophyll content only influences the light extinction ($K_h$) coefficient of leaf; **[b]** assumes low chlorophyll content not only affect $K_h$, but also $V_{cmax}$ and $J_{max}$ of leaf (Walker er al., 2018).

B. Gross $CO_2$ assimilation of pod.

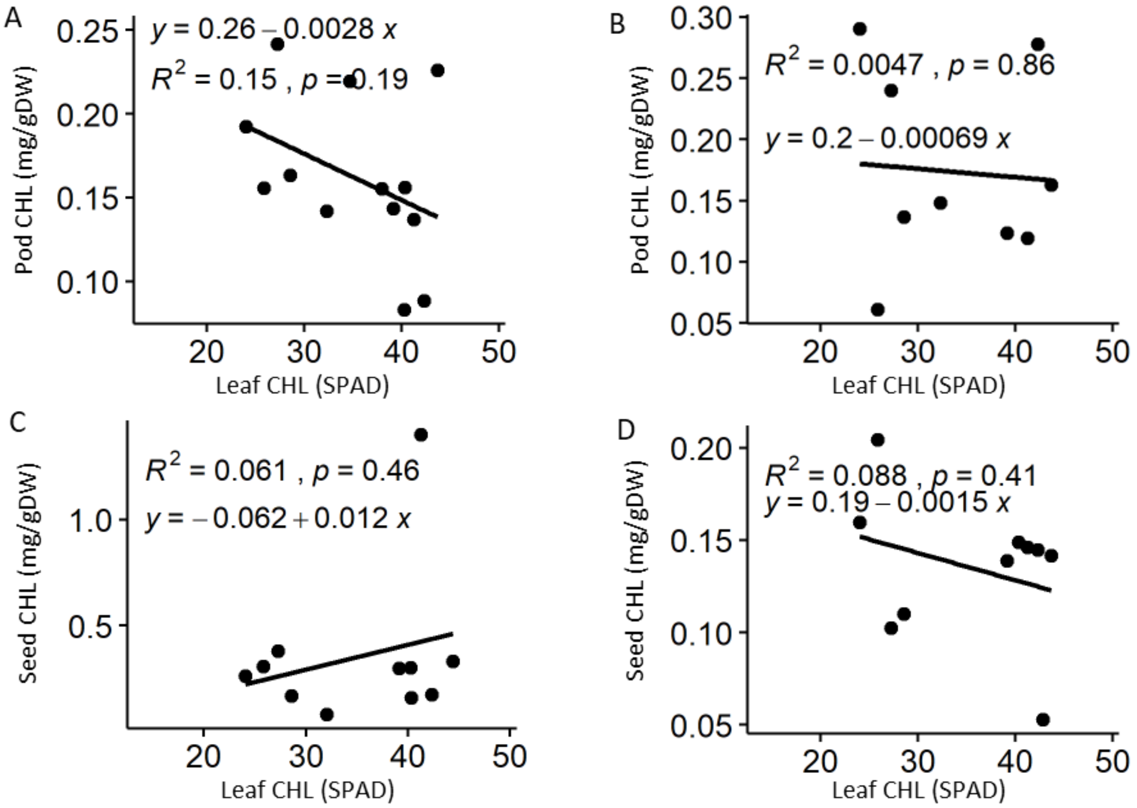


**Supplemental Figure 14 (Field). Correlation between the level of leaf chlorophyll (x-axis: SPAD reading) and the level of immature pod or seed chlorophyll (y-axis, mg/g DW).**

Line represents the linear regression model. R-squared is a coefficient of determination, the percentage of the response variable variation that is explained by the linear model. Pod is labeled by the fresh weight of seeds it contained.

A. Level of chlorophyll of 25-100mg pod (n=13).

B. Level of chlorophyll of 100-200mg pod (n=9).

C. Level of chlorophyll of 25-100mg seed (n=11).

D. Level of chlorophyll of 100-200mg seed (n=10).

| Variety | Type | Family | Chlorophyll concentration (mg/g dry weight) | | | | | Ratio (Leaf/tissue) | | | |
|---|---|---|---|---|---|---|---|---|---|---|---|
| | | | Leaf | Pod, 25-100 mg | Pod, 100-200 mg | Seed, 25-100 mg | Seed, 100-200 mg | Pod, 25-100 mg | Pod, 100-200 mg | Seed, 25-100 mg | Seed, 100-200 mg |
| 548217 | Mutant | Clark | 7.19 | 0.19 | 0.29 | 0.26 | 0.16 | 37.40 | 24.78 | 27.33 | 45.01 |
| 547555 | Mutant | Clark | 4.38 | 0.16 | 0.06 | 0.31 | 0.20 | 28.09 | 71.86 | 14.23 | 21.42 |
| 548265 | Mutant | Williams | 5.13 | 0.24 | 0.24 | 0.38 | 0.10 | 21.21 | 21.39 | 13.47 | 49.99 |
| 548289 | Mutant | Williams | 5.46 | 0.16 | 0.14 | 0.17 | 0.11 | 33.43 | 39.89 | 32.20 | 49.53 |
| 548215 | Mutant | Hawkeye | 8.39 | 0.14 | 0.12 | 0.30 | 0.14 | 58.51 | 67.83 | 28.01 | 60.45 |
| 548533 | Parent | Clark | 9.04 | 0.14 | 0.12 | 1.40 | 0.15 | 65.98 | 75.69 | 6.45 | 61.78 |
| 548362 | Parent | Lincoln | 9.37 | 0.09 | 0.28 | 0.17 | 0.14 | 106.15 | 33.77 | 53.96 | 64.68 |

**Supplemental Table 1 (Field). Comparison of chlorophyll level between leaf, pod and seed tissues.** Leaf chlorophyll level converted from SPAD values. Pods categorized by the fresh weight of seeds they contained.

| Variety | Type | Family | Chlorophyll concentration (mg/g dry weight) | | | | | Ratio (Leaf/tissue) | | | |
|---|---|---|---|---|---|---|---|---|---|---|---|
| | | | Leaf | Pod, 25-100 mg | Pod, 100-200 mg | Seed, 25-100 mg | Seed, 100-200 mg | Pod, 25-100 mg | Pod, 100-200 mg | Seed, 25-100 mg | Seed, 100-200 mg |
| 547555 | Mutant | Clark | 4.38 | N/A | 0.62 | 0.82 | 0.34 | N/A | 7.04 | 5.31 | 13.01 |
| 548231 | Mutant | Clark | 5.12 | 0.63 | 0.42 | 0.63 | 0.97 | 8.09 | 12.31 | 8.07 | 5.25 |
| 548265 | Mutant | Williams | 5.13 | 0.87 | 0.60 | 0.75 | 0.50 | 5.90 | 8.56 | 6.83 | 10.34 |
| 548204 | Mutant | Lincoln | 5.93 | 0.61 | 0.38 | 0.69 | 0.57 | 9.76 | 15.66 | 8.54 | 10.47 |
| 547452 | Mutant | Clark | 6.36 | 0.25 | 0.25 | 0.42 | 0.28 | 24.98 | 25.02 | 14.98 | 22.34 |
| 548206 | Mutant | Richland | 8.71 | 0.42 | 0.55 | 1.03 | 0.92 | 20.57 | 15.81 | 8.43 | 9.50 |
| 548402 | Parent | Peking | 8.74 | 0.57 | 0.40 | 1.07 | 0.60 | 15.31 | 21.59 | 8.20 | 14.51 |
| 548533 | Parent | Clark | 9.04 | 0.51 | 0.39 | 1.04 | 0.78 | 17.71 | 23.45 | 8.69 | 11.59 |
| 548221 | Parent | T239 | 9.69 | 0.98 | 0.70 | 1.09 | 0.76 | 9.92 | 13.81 | 8.86 | 12.78 |
| 540553 | Parent | T322 | 9.81 | 0.92 | 0.67 | 0.86 | 0.80 | 10.70 | 14.59 | 11.35 | 12.24 |

**Supplemental Table 2 (greenhouse). Comparison of chlorophyll level between leaf, pod and seed tissues.** Leaf chlorophyll level converted from SPAD values. Pods categorized by the fresh weight of seeds they contained.
